# Causal Discovery Analysis Reveals Insights into Psychosis Proneness, Brain Function, and Environmental Factors among Young Individuals

**DOI:** 10.1101/2024.02.19.581044

**Authors:** Tuba Sahin-Ilikoglu, Sisi Ma, Erich Kummerfeld, Eric Rawls, Hao Yang Tan, Timothea Toulopoulou

## Abstract

**Background:** Experiencing symptoms of psychosis, such as delusions and hallucinations, are observed in general, nonclinical populations. These experiences are sometimes described as psychosis proneness (PP) and potentially part of the psychosis continuum. The directional relationships among various factors contributing to psychosis proneness and its interactions, encompassing both environmental and neural mechanisms, lack comprehensive description. We aimed to identify targets to prevent psychosis proneness and its interactions by characterizing pathways using causal discovery analysis (CDA).

**Methods:** Participants were 194 healthy adolescent and young adult twin and sibling pairs aged between 14-24 years from Türkiye. They completed comprehensive assessments evaluating sociodemographic status, environmental risk, general intelligence, self-schema, PP, and working memory (WM) performance during fMRI (37 variables). CDA was applied, a novel machine learning algorithm, to understand the causal relationships of PP.

**Results:** The analysis identified negative self-schema as having the largest causal effect among all assessments in PP [Effect size (ES)= 0.55]. Secondly, social cohesion and trust (SC&T) had a protective causal effect on PP [ES= −0.18]. Lastly, PP was identified as a direct cause of greater activation in DLPFC (BA9a-BA46v) during manipulation in the WM (ES= 0.14).

**Conclusions:** CDA provides directionality of the 37 variables which were not presented earlier. The findings highlight the significance of negative self-schema and SC&T in the general population with PP, emphasizing the potential for preventive interventions targeting these factors. These findings also suggest a role for DLPFC as a potential target in this regard. To our knowledge, this is the first study using data-driven analysis to model causal mechanisms in PP in the general population.

## INTRODUCTION

Psychotic-like experiences are also reported in the general population, particularly during adolescence and young adulthood (Kelleher et al., 2012; van Os, Linscott, Myin-Germeys, Delespaul, & Krabbendam, 2009). Understanding what gives rise to these symptoms might be important as their presence during adolescence correlates with an increased risk of developing psychotic (Poulton et al., 2000) and other psychiatric conditions (Bourgin et al., 2020) later in life.

Previous research has indicated that environmental adversity not only elevates the risk of developing schizophrenia but also influences the lower end of the psychosis spectrum, such as phenotypic psychotic proneness in healthy populations and unaffected siblings (Varese et al., 2012). Various environmental risk factors for psychosis have been identified, including obstetric complications (Clarke & Cannon, 2020), migration, minority group position (Bourque, van der Ven, & Malla, 2011), urban birth and urban upbringing (Krabbendam & van Os, 2005), childhood trauma (Varese et al., 2012), and cannabis use (Arseneault et al., 2002). These factors have been associated with neural disruptions, leading to a greater vulnerability and alterations in brain structure and function, especially in the prefrontal cortex (Teicher, Samson, Anderson, & Ohashi, 2016; van Os, Kenis, & Rutten, 2010).

Alongside environmental factors, cognitive deficits have emerged as a crucial feature of psychotic disorders. These deficits, commonly found among healthy relatives of people with schizophrenia (Goldberg et al., 1995; Toulopoulou, Rabe-Hesketh, King, Murray, & Morris, 2003) are thought to precede psychotic symptoms (Reichenberg et al., 2010) and may play a causal role (Toulopoulou et al., 2015). Cognitive dysfunction may, therefore, serve as an early indicator of individuals vulnerable to psychosis before the full onset of the disease.

Moreover, disruptions in the cognitive framework that shapes beliefs about oneself (self-schema), have frequently been reported in psychotic disorders. For example, negative self-perception and negative self-concepts have been found to be associated with paranoia (Palmier-Claus, Dunn, Drake, & Lewis, 2011), hallucinations (Aaron T. Beck & Rector, 2003), and delusions (Freeman, Garety, & Kuipers, 2001) and may interact with environmental factors (Garety, Bebbington, Fowler, Freeman, & Kuipers, 2007), although the nature and direction of this interaction remains unclear. Research exploring PP in relation to self-perception is limited.

Despite progress in understanding the mechanisms that underlie PP and its interactions, comprehensive assessments are lacking. Previous research investigating PP largely employs traditional statistical methods, which fall short of capturing potential causal factors. To address this gap, we took a comprehensive approach to assessment spanning environmental, cognitive, clinical, and neurobiological data and used causal discovery analysis, a novel method that differentiates causation from association to identify causal pathways (cause and effect) to PP and its interactions in the general population. Unlike expert-based structural equation modeling (SEM), which is dependent on a priori expert opinion, CDA is data-driven, albeit with the option to incorporate expert background knowledge. In SEM, it is presupposed that no variables are missing, and the correct model is pre-specified, often requiring the reduction of variables for accurate modeling. Conversely, CDA introduces all relevant variables, allowing algorithms to identify the best-fitting model without the need for a priori assumptions or variable reduction (Spirtes, Glymour, & Scheines, 2000). Notably, SEM typically relies on theory and necessitates the a priori specification of latent variables, whereas Greedy Fast Causal Inference (GFCI) searches for causal relationships directly within observed data, avoiding explicit modeling of latent variables (Ogarrio, Spirtes, & Ramsey, 2016). This distinction underscores SEM’s focus on confirming a predefined model, while GFCI is tailored for inferring causal structures from observational data. Since our approach was data-driven, while we anticipated discovering connections between PP, environmental risk, negative self-schema, and neurobiological deviations, we were mostly agnostic about the direction of causation.

## METHODS

### Participants

Participants were from a larger study on Brain Development and Psychosis focusing on a healthy adolescent and young twin and sibling population (Tubitak 1001, 119K410, PI TT). The current study included data from 194 participants, collected between 20.04.2017 and 11.08.2022. Inclusion/exclusion criteria are described in Supplementary. Participants were aged between 14-24 (M= 19.8, SD=2.34) and were Turkish citizens with the Turkish mother tongue. All participants gave written consent before participation. A legal guardian gave consent to participants under the age of 18. The study was approved by the Human Research Ethics Committee of Ihsan Dogramaci Bilkent University and Ankara University Human Ethics Committee.

### Psychological Assessments

#### Community Assessment of Psychic Experiences (CAPE-42)

PP was assessed using the CAPE-42. CAPE-42 is a 42 items self-report questionnaire that measures lifetime psychotic experiences and PP in the affective and non-affective domains (Stefanis et al., 2002). CAPE-42 measures frequency and distress related to psychotic experiences, and there are three subscales: positive, negative and depression. Meta-analysis on the psychometric properties of CAPE-42 showed that it is a reliable and valid questionnaire, especially among the young population, to measure PP (Mark & Toulopoulou, 2015). For the study, only total CAPE-42 score was used.

#### Brief Core Schema Scales (BCSS)

BCSS (Fowler et al., 2006) is a 24-item self-report assessment of schemata concerning self and others (see supplementary). Extreme negative evaluations of self and others appear to be characteristic of the appraisals of people with chronic psychosis and are associated with symptoms of grandiosity and paranoia in the non-clinical population.

#### Environmental Assessments

Ten measures of well-established environmental risk factors for psychosis were included: Urbanicity, Obstetric Complications, Cannabis Use, Childhood Trauma, Parental Discord Rating Scale, Alcohol Usage, Discrimination, Life Threatening Events, Socia Cohesion and Trust (SC&T). Please see supplementary methods for details and scoring of all variables.

#### Family Interview for Genetic Studies (FIGS)

The FIGS was used to assess whether participants have first-degree relatives (parents, siblings, children) with psychosis and schizophrenia as a structured interview (“NIMH Genetics Initiative: Family Interview for Genetic Studies (FIGS),” 1992).

## COGNITIVE ASSESSMENTS

**Wechsler Abbreviated Scale of Intelligence | Second Edition (WASI-II).** Block-Design and Matrix-Reasoning from the Revised Wechsler Abbreviated Scale of Intelligence (WASI-II) was used as a quick measure of intelligence (Wechsler, 2011) (see supplementary). **Verbal Fluency.** Verbal fluency was measured in two categories: letter fluency (Turkish version) and category fluency using standard assessment and scoring criteria (see supplementary) (Borkowski, Benton, & Spreen, 1967). Participants were asked to name as many words as possible during a period of 60 seconds (Borkowski et al., 1967).

## NEUROIMAGING

Information on MRI apparatus, task-based functional magnetic resonance imaging (fMRI) and image preprocessing can be found in the online supplement.

### Working Memory Paradigm

We used an event-related paradigm that previous studies in patients with schizophrenia and their healthy first-degree relatives showed alterations in cortical activations and behavioral performance (Greenman et al., 2020; Tan et al., 2007) particularly during WM manipulation-the focus of the fMRI analysis in this paper. Thus, the task is sensitive in capturing the neural and cognitive deviations in both patients and healthy individuals who are vulnerable to psychosis. The paradigm has been described in detail elsewhere (Tan et al., 2007) and see supplementary for details.

## DATA ANALYSIS

### fMRI Analysis

#### Statistical Analysis

General Linear Model (GLM) specification was modeled using the canonical hemodynamic response function (HRF). The ratio was normalized to the whole-brain global mean to control for systematic differences in global activity and temporally filtered with a high-pass filter of 128 s. Events were modeled for both correct and incorrect trials of event-related designated cognitive paradigm during each task phase. In addition, the residual movement and incorrect response parameters were modeled as regressors of no interest. Cognitive subtraction was performed for “Manipulation” by subtracting Motor task (M) from Compute/estimate after encoding (E_CJ) (Fig 1; Supplementary Fig 1).

**Figure 1.**
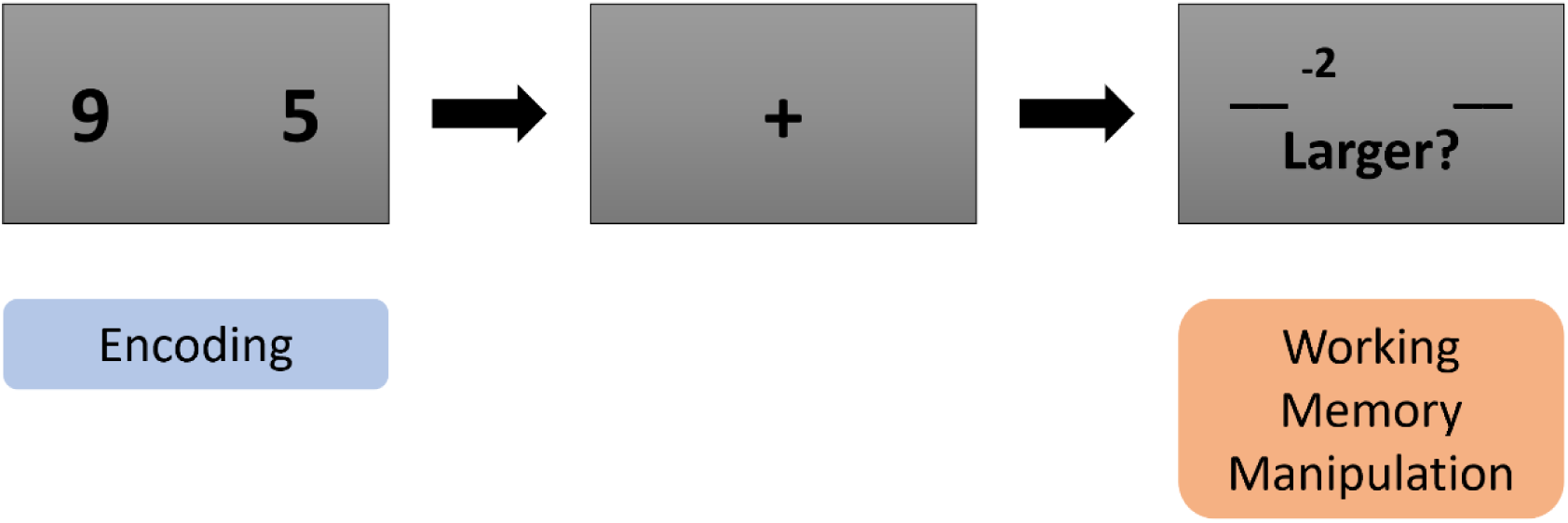
Working Memory Paradigm. During encoding phase, two number digits were presented in the left and right side of the screen. The location and the digit numbers were held in working memory for 3 to 6 seconds. In the working memory manipulation phase, subjects performed subtraction on the retrieved number of the indicated side in working memory, and participants were asked to respond which side of the number was larger or smaller.

#### Regions of Interest (ROI) Analysis

ROIs were defined for the task-based fMRI during the manipulation phase of WM. Peak activations were identified for 16 regions (Fig 2 & Supplementary Table 1) by using the Human Connectome Project MMP extended Atlas, HCPex (Huang, Rolls, Feng, & Lin, 2022) in MRIcron. Only ROIs that passed FWE-correction at peak level were included.

**Figure 2.**
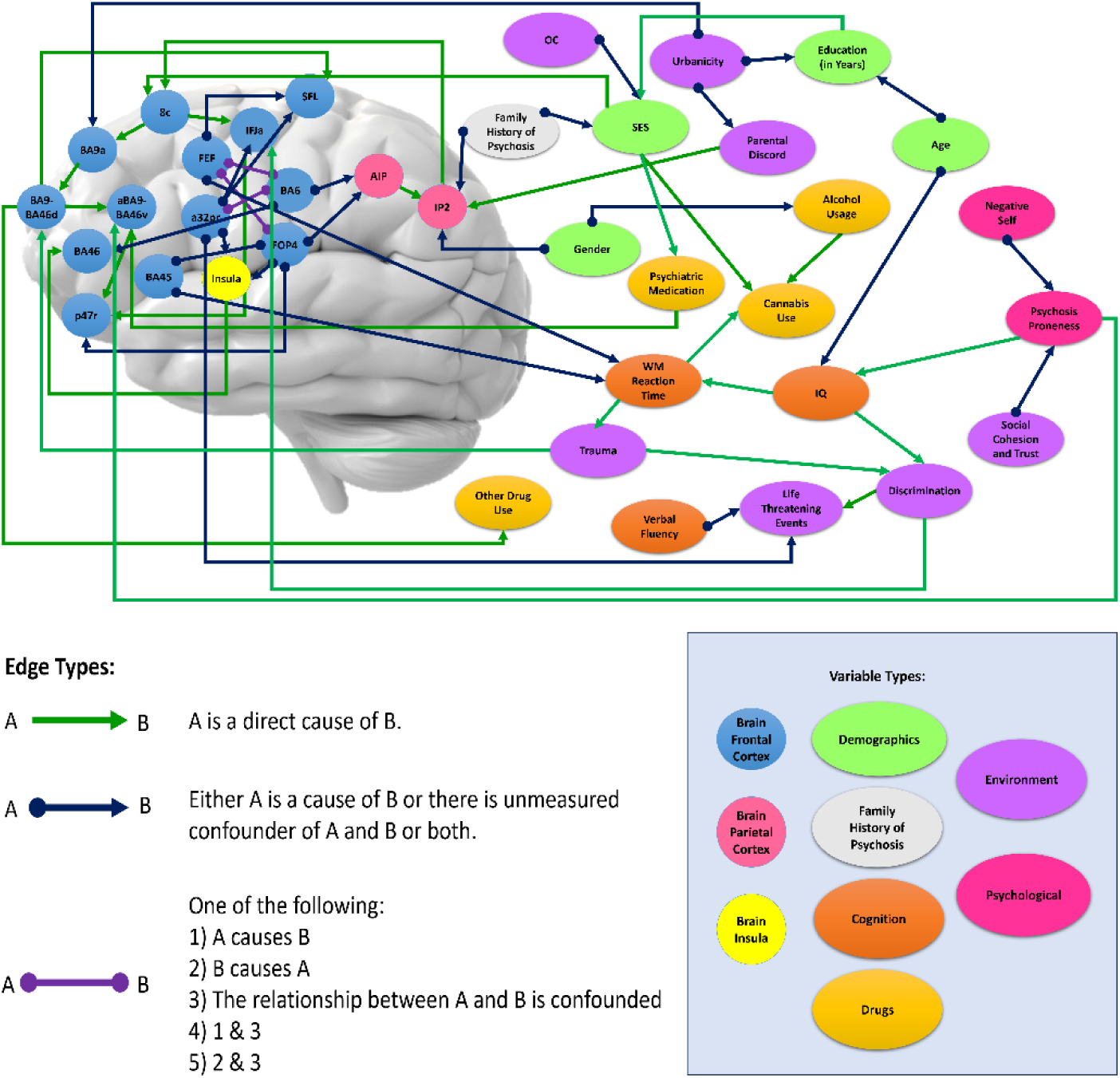
Causal Discovery Analysis for all variables and Manipulation phase of the Working Memory Paradigm. Causal Discovery Analysis was performed using GFCI for activations in ROIs when manipulating information during fMRI, environmental and psychological variables. (B.A.:Brodmann; SFL: Superior frontal language area; AIP: anterior intraparietal sulcus, I.P.: Intraparietal Sulcus; a9-46v: frontal mid, FOP4: frontal operculum, IFJa: anterior inferior frontal gyrus, p47r: the orbital prefrontal cortex, a32pr: refers to a narrow strip of the anterior midcingulate cortex, FEF: frontal eye fields; O.C.: obstetric complications; SES: socioeconomic status; W.M.: working memory) (Brain template was taken with permission from https://www.mindmedia.com/en/solutions/qeeg/brain-mapping/).

#### Causal Discovery with Greedy Fast Causal Inference

The observed variables were adjusted for family-level clustering to ensure that observations were independent of family effects (see supplementary). Data-driven causal modeling was applied to examine the relationship between variables presented in Table 1 and ROIs (Supplementary Table 2). In total 37 variables were introduced to CDA. Causal Discovery Analysis (CDA) was performed with the Greedy Fast Causal Inference (GFCI) algorithm (Ogarrio et al., 2016) implemented in the Tetrad software package version 6.9 (https://cmu-phil.github.io/tetrad/manual/). GFCI combines the Greedy Equivalence Search (GES) algorithm and the Fast Causal Inference (FCI) algorithm. The procedure has been described in detail before (Miley et al., 2023). Briefly, it applies GES to find a causal skeleton followed by FCI to test for errors caused by unmeasured common factors in relationships (confounders). GES algorithm initially starts its Causal Bayesian Networks search with an empty graph with no edges between variables. Afterwards, GES adds edges between nodes (variables) one at a time until a better fit model was reached for the observed data which means that adding or removing any single edge would not improve the overall fit. GFCI returns a partial ancestral graph (PAG) depicting causal relationships between a set of variables (PAG). In the generated PAG, nodes represent variables and edges represent modeled causal relationships. The best-fitting model during the GES step is assessed using the Bayesian Information Criterion (BIC) (Schwarz, 1978) with a penalty discount of 1 (that is, the standard BIC score). Utilizing the Bayesian Information Criterion (BIC) in this CDA is another notable aspect, as it plays a role akin to multiple comparison corrections, penalizing intricate models by pruning connections. This mechanism aids in curbing false associations, particularly those that are weaker in nature. In the second phase of the GFCI algorithm, which is FCI, starts with an undirected graph generated during the GES algorithm. FCI algorithm checks the pairs of variables that are adjacent and checks for conditional independence relations between pair of variables. If the condition set between a pair of variables is found to be independent, the edge between them is removed. Conditional independence tests during the FCI step uses Fisher’s Z test, with alpha set to the default value of 0.01. Then FCI tries to find the direction of the remaining edges. At this final step, direction of edges is not only searched with conditional independence test performed during FCI but also the information from previous step, GES, is used. GFCI finds latent variables via the patterns of conditional independence. GFCI considers the possibility of latent variables when certain variables become independent given another set of variables during the conditional independence. We used GFCI to develop hypotheses about putative causal pathways between psychosis proneness, self-schema, environmental risk factors, cognitive factors, demographics, and fMRI variables. We used the default parameterization for GFCI. The GFCI algorithm can discover causal relationships and potential presence of hidden common causes among variables up to statistical equivalence under broad assumptions. The output of the GFCI algorithm is a partial ancestral graph (PAG). See Supplementary for more detailed information on GFCI. The stability of the generated PAG was evaluated by applying GFCI to resamples of the original dataset at a 90% resampling rate, repeated 1000 times.

**Table 1.** Sample Characteristics.

| **Sample Characteristics** | **N=194** | **Min** | **Max** |
| --- | --- | --- | --- |
| **Demographic Variables** |  |  |  |
| **Age, mean (SD)** | 19.8 (2.34) | 14 | 24 |
| **Gender** |  |  |  |
| *Female, n (%)* | 80 (40.82%) |  |  |
| *Male, n (%)* | 116 (59.18%) |  |  |
| **Sibling Type** |  |  |  |
| *M.Z, n (%)* | 64 (32.9%) |  |  |
| *D.Z., n (%)* | 98 (50.52%) |  |  |
| *Sibling, n (%)* | 32 (16.9%) |  |  |
| **Socioeconomic status of mother or father** |  |  |  |
| *have never worked or have been unemployed for a long time* | 2 (1.03 %) |  |  |
| *routine work* | 4 (2.06 %) |  |  |
| *lower-level technical job* | 18 (9.28 %) |  |  |
| *lower-level service, sales, secretarial* | 14 (7.22 %) |  |  |
| *lower-level administrative or technician* | 12 (6.19 %) |  |  |
| *self-employed* | 25 (12.89 %) |  |  |
| *small medium business owner* | 12 (6.19 %) |  |  |
| *intermediate level* | 67 (34.54 %) |  |  |
| *middle level manager* | 32 (16.49 %) |  |  |
| *senior manager* | 8 (4.12 %) |  |  |
| **Years of Education, mean (SD)** | 13.89 (2.12) | 9 | 18 |
| **Psychological Variables** |  |  |  |
| Psychosis Proneness, mean (SD) | 74.32 (15.4) | 42 | 122 |
| Negative Self-Schema, mean (SD) | 5.33 (4.74) | 0 | 22 |
| **Environmental Variables** |  |  |  |
| **Ethnic Minority** |  |  |  |
| *Turkish, n (%)* | 170 (87.63 %) |  |  |
| *Other, n (%)* | 24 (12.37 %) |  |  |
| **Paternal Age** |  |  |  |
| <40, n (%) | 174 (89.69 %) |  |  |
| 40-50, n (%) | 18 (9.28 %) |  |  |
| >50, n (%) | 2 (1.03 %) |  |  |
| **Obstetric Complications, mean (SD)** | 0.52 (0.98) | -.5 | 2.5 |
| **Parental Discord** |  |  |  |
| *None, n (%)* | 74 (38.14 %) |  |  |
| *Some, n (%)* | 15 (7.73 %) |  |  |
| *Moderate, n (%)* | 45 (23.20 %) |  |  |
| *Marked, n (%)* | 42 (21.65 %) |  |  |
| *Violence, n (%)* | 18 (9.28 %) |  |  |
| **Urban Birth** |  |  |  |
| *Urban center, n (%)* | 139 (71.65 %) |  |  |
| *Urban Cluster, n (%)* | 48 (24.74 %) |  |  |
| *Rural, n (%)* | 7 (3.61 %) |  |  |
| **Childhood Trauma** |  |  |  |
| *Trauma, n (%)* | 51 (26.29 %) |  |  |
| *No Trauma, n (%)* | 143 (73.71 %) |  |  |
| **Discrimination, mean (SD)** | 1.42 (1.67) | 0 | 8 |
| **Life-threatening events (Last 12 months), mean (SD)** | 2.52 (1.9) | 0 | 8 |
| **Social Cohesion and Trust, mean (SD)** | 30.1 (8.6) | 0 | 50 |
| **Alcohol Usage** |  |  |  |
| *>12 drinks in the last 12 months, n (%)* | 100 (51.55 %) |  |  |
| *<12 drinks in the last 12 months, n (%)* | 94 (48.45 %) |  |  |
| **Cannabis Use** |  |  |  |
| *No exposure, n (%)* | 147 (75.77 %) |  |  |
| *Little or a few times a year = Little/Moderate, n (%)* | 33 (17.01%) |  |  |
| *A few times a month & above = High Exposure, n (%)* | 14 (7.22 %) |  |  |
| **Drug usage other than cannabis** |  |  |  |
| *Used, n (%)* | 14 (7.22 %) |  |  |
| *Not used, n (%)* | 180 (92.78 %) |  |  |
| **Psychiatric Medication** |  |  |  |
| *Used, n (%)* | 31 (15.98 %) |  |  |
| *Not used, n (%)* | 163 (84.02 %) |  |  |
| **Family History of Psychosis** |  |  |  |
| *Existence, n (%)* | 12 (6.19 %) |  |  |
| *No Existence, n (%)* | 182 (93.81 %) |  |  |
| **Cognitive Variables** |  |  |  |
| IQ | 102.39 (13.65) | 72 | 154 |
| Working-memory reaction time, mean (SD) | 0.95 (0.3) | 0.19 | 1.92 |
| Verbal Fluency, mean (SD) | 13.09 (3.41) | 4.25 | 24.5 |

#### Causal Effect Size Estimation

We used the R package Lavaan 0.6.9 (Rosseel, 2012) to fit a SEM to the discovered PAG, to recover standardized effect sizes for each modeled causal pathway. Directed causal relationships were modeled as linear regressions and uncertain paths or otherwise undirected paths were modeled as covariances. Uncertain paths linked to psychosis proneness (negative self-schema and SC&T) were additionally modeled as direct paths in an additional SEM analysis. The interpretation of the causal effect size is the estimated change in the effect when the cause is changed by 1 standard deviation while holding all other variables constant. The model fit was examined using the Tucker-Lewis Index (TLI) and Root Mean Square Error of approximation (RMSEA).

## RESULTS

### Descriptive Statistics

Demographic and PP variables are reported in Table 1.

### Causal Discovery Analysis

GFCI was used to form a model for PP encompassing sociodemographic (4), environmental (7), psychological, including the PP (2), cognitive (3), drug usage (4), family history of psychosis (1), and fMRI indexes (16) during the manipulation of information in WM (Fig 1). The output of GFCI is shown in Fig 2 and Supplementary Fig 2 with effect size and information on p-values.

The SEM fit to this model indicated a good fit, RMSEA= 0.052, Tucker-Lewis Index = 0.833. Edge weights extracted from SEM are presented next to the corresponding arrow on the graph in Fig 3. Except for four edges, all paths are significant at p < .05, and 39 out of 57 at p < .01 (Fig 3, Supplementary Fig 2). As standard, we used uncorrected p-values on graphs. The results from the Benjamini Hochberg method to control False Discovery Rate (FDR) are in supplementary Table 2. After FDR correction, 53 edges out of 57 survived at p < 0.05. We found no difference in statistical significance between uncorrected p values and FDR corrected p values. (Supplementary Table 2).

**Figure 3.**
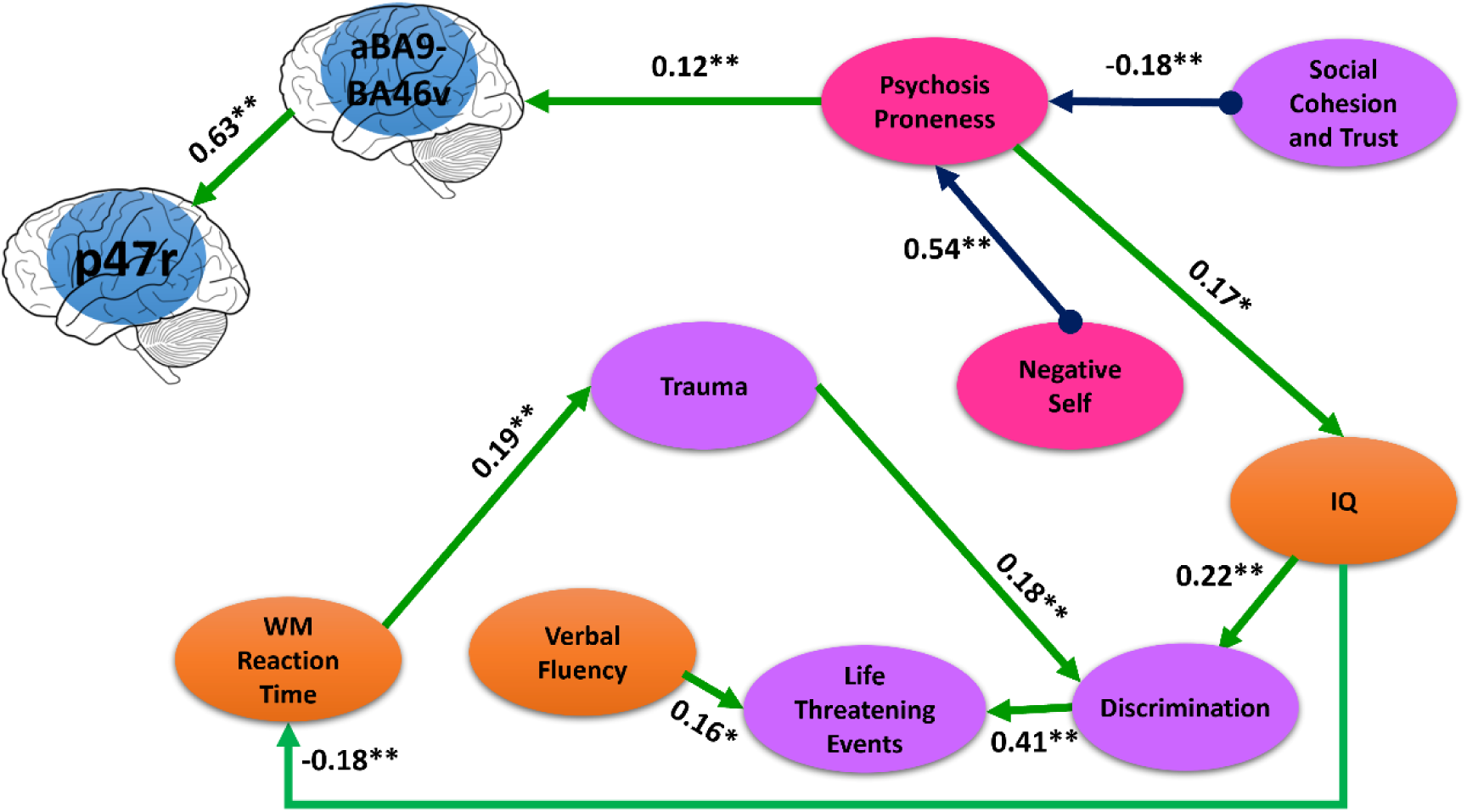
Psychosis Proneness Subgraph. GFCI returns a partial ancestral graph (PAG) depicting causal relationships between a set of variables, while assessing for unmeasured third variables in relationships (confounders). BA9: DLPFC; BA46: MPFC; P47r: Orbital Prefrontal Cortex

### Causal Model Results Interpretation

The causal model graph (Fig 2) illustrated a complex network of links between introduced variables. While some relationships are direct, others exhibit complex dynamics, potentially involving latent variables that might be overlooked with human-driven hypotheses. The relationships involving the possibility of a latent variable are referred to as “possibly confounded” relationships, that a possibility of a latent factor should be considered (see supplementary).

One of the notable findings from the CDA is the positive and statistically significant possibly confounded relationship between the negative self-schema and PP (ES = 0.540, p < 0.001). The obtained result indicates that higher levels of negative self-schema is associated with increased PP. We also found influences of SC&T on PP (ES= −0.18, p=0.003), indicating that increased levels of sense of belonging and trust towards the social environment lead to decreased levels of PP. Importantly, there was a latent (unobserved) factor between the negative self-schema and PP and SC&T and PP, implying that some unmeasured variable might be a common cause of both negative self-schema and PP and SC&T and PP. Furthermore, the relationship between PP to IQ was found (SE= 0.17, p < 0.05). Our result might indicate that people prone to psychosis tend to have higher level of IQ.

We further found that PP influenced activation in the BA9-BA46v, a part of the dorsolateral prefrontal cortex (DLPFC). The analysis revealed a strong and significant positive relationship (ES = 0.12, p = 0.0013). Furthermore, aBA9-BA46v activation exhibited a positive effect on activation in p47r (ES = 0.63, p < 0.001).

The graph suggested the possibly confounded statistically significant (p< 0.01) causal effect of the mid ventrolateral prefrontal cortex (VLPFC), (∼B.A. 45; inferior frontal gyrus pars triangularis) to WM RT (ES=-0.17). This result indicates that the higher activation in VLPFC causes lower RT during WM performance.

The graph suggested other diverse causal relationships of cognitive variables with other variables. For example, we observed the causal effect of IQ to WM RT (ES= −0.18, p < 0.01), which suggests higher IQ is related to lower WM RTs, thus faster responses. We also observed positive causal relationship between WM RT and trauma (ES= 0.19, p <0.05). Further, trauma had causal effect on discrimination (ES= 0.2, p<0.01) and discrimination had causal effect on life-threatening events (ES= 0.41, p <0.01). The latter, in turn, was influenced by an unmeasured factor that also influenced verbal fluency (ES= 0.16, p <0.5). Finally, a possibly confounded causal effect of urban birth of the child on parental discord severity (ES= −0.2, p < 0.5) was observed.

We identified several significant causal relationships between fMRI variables and other variables, distinct from PP. Specifically, effect of trauma to BA9-BA46d (ES= −0.12, p < 0.5) was found, while parental discord exhibited negative effect to IP2 (ES= 0.11, p < 0.05). Specifically, a32pr activation was found to have a possibly confounded negative effect on the occurrence of life-threatening events (ES = −0.12, p < 0.05), while psychiatric medication exhibited a negative effect on DLPFC (ES= −0.1, p < 0.01). Additionally, BA9-BA46d activation was associated with drug usage, excluding cannabis (ES = −0.18, p < 0.01), and SES showed a positive effect on 8c activation (ES = 0.27, p < 0.001). A possibly confounded relationship was observed between family history of psychosis and activation in IP2 (ES = −0.1, p < 0.05). Furthermore, a possibly confounded relationship between Gender and IP2 was observed (ES= −0.1, p < 0.05).

## DISCUSSION

In this study, we utilized CDA, a data-driven machine learning approach, to characterize causal transactional pathways linking PP to multimodal factors, including cognitive variables, self-schema, environmental factors, and functional brain activation in WM, a cognitive process particularly affected in schizophrenia and their first-degree relatives.

In our study, we examined the complex nature of PP, revealing both its causal origins and consequences within the sample. Among the various factors we explored, negative self-schema emerged as a possibly powerful contributor, significantly influencing the manifestation of PP with a substantial effect size of 0.55. Moreover, SC&T was also possibly significantly influencing the PP. Furthermore, our findings unveiled a neurobiological insight, demonstrating that there is a direct influence of PP on the increase in activation within the DLPFC during WM.

Our main finding is that negative self-schema has a possibly confounded causal role in PP. This finding is in line with other studies showing that patients with psychosis, including people at risk, have maladaptive self-schemas (Fowler et al., 2006; Palmier-Claus et al., 2011) that often start early such as in childhood and adolescence. Our study extends these earlier findings in people with PP while utilizing an extensive range of cognitive, environmental, and neurobiological assessments in addition to an assessment of self-schema. Our results showed that negative self-schema was the possibly strongest cause contributing to PP compared to other well-established risk factors of psychosis. Self-disturbances in schizophrenia often comprise depersonalization, derealization, thought and perceptual disturbances regarding the understanding of the self and others, leading to a fragile sense of self and alienation from others (Raballo, Poletti, Preti, & Parnas, 2021). Although our study did not support the role of early trauma in PP, others have reported early traumatic experiences affect cognition and cause negative perceptions of self and others (A. T. Beck, 2008). The identified causal relationship between negative self-schema and PP provides valuable insights into the potential psychological mechanisms underlying the development of psychosis-related symptoms. Our results suggest that negative self-schema might serve as a filtering system for PP. Individuals who exhibit more negative self-attributes may be at a greater risk of experiencing symptoms related to psychosis.

Previous studies had shown the protective effect of social cohesion by interacting with stressful life events on mental health of adolescents such as anxiety and depression (Kingsbury, Clayborne, Colman, & Kirkbride, 2020). We found significantly possibly direct causal relationship between SC&T and PP which suggests that higher levels of SC&T possibly leads lower levels of PP. Therefore, our result suggests the role of SC&T on not only during disease but also for its effect on PP. Additionally, a direct effect of urban birth on PP was not observed, while an effect of SC&T was observed, suggesting different roles in vulnerability to psychosis.

PP has a direct causal relationship to DLPFC in our sample. This finding aligns with previous studies that have documented alterations in neuronal activity among participants who subsequently developed psychosis, thereby bolstering our conclusion (Tan, Choo, Fones, & Chee, 2005). The effect size of 0.12 from PP to neural activity in DLPFC is low but statistically significant. Initially, psychosis proneness effect on functional activity in brain involves two different dimensions of measurements. While psychosis proneness was measured by psychometric data, the latter involves BOLD signal during working memory performance. DLPFC plays a role in executive functions including WM, which are important for daily functioning (Curtis & D’Esposito, 2003). The result highlights a potential neural substrate associated with PP. The significant relationship suggests that increased levels of PP lead to altered neural activity in the DLPFC during manipulation of information in WM tasks. Such findings may indicate a link between cognitive deficits and PP. WM is disturbed in schizophrenia (Meyer-Lindenberg et al., 2001) and in psychosis prone people (Colibazzi et al., 2016; Wolf et al., 2015). Given that full-blown psychosis and PLEs share the same continuum, it is not unexpected that PP has a causal role on DLPFC during WM performance. Our earlier work also suggested a causal role of schizophrenia liability to brain changes (Toulopoulou et al., 2015). In our study, increasing PP resulted in increased activation in DLPFC. This finding in line with the literature that control samples perform better during manipulation in WM and therefore associated with increased activation in DLPFC compared to patients (Tan et al., 2005).

In contrast to the prevailing findings that individuals with schizophrenia tend to have lower IQ levels (Maccabe, 2008), our current data suggests a potential link between PP and higher IQ. Some subgroups within the schizophrenia patient population also show higher IQ levels (MacCabe et al., 2012) and exhibit distinct symptom profiles. Additionally, exceptional intellectual abilities during adolescence have been linked to later development of bipolar disorder (MacCabe et al., 2010). However, research on the relationship between PP and IQ in the general population is limited, preventing firm conclusions.

Earlier studies have shown that both DLPFC and VLPFC play a role in aberrant activation during WM (MacDonald, Thermenos, Barch, & Seidman, 2009). Although we did not observe a relationship between WM behavioral performance (RT) with DLPFC, we did observe a confounded causal relationship with the VLPFC. Furthermore, we also found a possibly confounded causal effect of family history of psychosis on IP2, which suggests the existence of a family history of psychosis results in the reduced activation of IP2. Since we could not incorporate molecular genetic data for schizophrenia into our model, the possibly confounded relationship between WM performance and VLPFC, and family history of psychosis and IP2 might be further explored.

Some of the findings derived from the data-driven approach are hard-to-explain results (for bivariate relations exclusive of PP such as Verbal fluency and Threatening events; Urbanicity and Parental Discord; Gender and activation in IP2; Age and IQ; activation in a32pr and life-threatening events). Future research is necessary to provide needed clarity on their causal relationships.

The absence of an association between some environmental risk factors, such as trauma, cannabis use, family history of psychosis and PP in our study could be attributed to two key factors. First, the relatively low presence of PP in our sample may have played a role in this outcome. Secondly, we did not employ a stratification method during recruitment, such as distinguishing between cannabis users and non-users, resulting in a limited frequency of individuals exposed to specific environmental risk factors. Our primary focus was not to investigate the effect of a single factor such as cannabis usage, but rather to observe the directionality of all available risk factors in the general sample. Furthermore, previous studies have typically examined cannabis use in relation to specific symptomatology, such as positive schizotypy, which may not provide a comprehensive representation of psychosis proneness (Radhakrishnan, Kamal, Guloksuz, & van Os, 2023; Szoke et al., 2014). Additionally, our lack of direct association between cannabis use and PP is in line with previous research that lifetime cannabis use was not statistically significantly associated with CAPE scores but with other psychometric measures (O’Tuathaigh et al., 2020). Consequently, we might not observe the direct effect of these factors on psychosis proneness. It’s worth noting that our participants did not exhibit symptoms of psychosis, which might have contributed to this lack of association. Future studies can investigate the effects of different measures of psychosis proneness.

These findings support the notion that PP is influenced by both negative self-schema and SC&T and impact brain activation patterns during WM task. The bidirectional causal relationships between brain mechanisms and PP suggest a complex interplay between cognitive control processes and the development of psychosis-related traits. Moreover, the observed effects of a decreased sense of belonging and trust on PP highlight the significance of social environmental factors in contributing to psychosis vulnerability. The positive relationship between PP and activation in the DLPFC suggests a potential neural basis for the manifestation of PP during cognitive tasks. The application of CDA yielded superior insights into the directionality of the relationships in comparison to the traditional analysis such as regression analysis. Notably, CDA’s automated methods like GFCI have performed well on real-world data (Shen et al., 2020) enhancing the credibility of this endeavor. By shedding light on the multifaceted connections between psychological and neurobiological factors, our research helps progress towards a deeper understanding of psychosis and offers valuable implications for future interventions and treatment approaches. To support the hypothesized causal pathways, an empirical strategy could encompass focused interventions designed to alter self-schemas and SC&T followed by an assessment of potential decreases in PP scores.

### Limitations and Future Directions

While the current study offers novel information on data-driven causal discovery for PP among the healthy populations, our results should be interpreted in light of limitations. We acknowledge the consideration of incorporating the genetic risk in aggregate such as polygenic risk score (PRS) into our data-driven approach. These scores can explain ∼ 9.9 % of variance in liability to schizophrenia and 4.3 % of variance in general cognitive function (Davies et al., 2018; Guloksuz et al., 2019; Trubetskoy et al., 2022). First degree family history of psychosis, which is a proxy of genetic risk, was included in the CDA but did not provide additional insights into the relationship. Given our general population sample with no participant diagnoses, the prevalence of first-degree diagnoses was relatively low (n=12). Existing research highlights a substantial heritability gap in schizophrenia, with Genome-wide Association Studies (GWAS) explaining only a small portion (Consortium, 2014; Guloksuz et al., 2019). Although introducing the PRS could address some concerns, we anticipate a diminished and less significant genetic role in our general population sample. Nonetheless, integrating the PRS factor may elucidate the confounded causal relationship between negative self-schema, social cohesion, and trust in psychosis proneness, emphasizing the continued importance of exploring environmental and other factors. Therefore, including these scores in future studies will enrich the understanding of the factors that contribute to PP. Furthermore, subsequent research, employing larger sample sizes, may further investigate potential confounded relationships, considering the continuum between vulnerability to psychosis proneness and depression/anxiety disorders (van Os & Reininghaus, 2016; Verdoux et al., 1999). Expanding on this, future studies could explore vulnerability to various mental disorders, as individuals with clinical depression often exhibit increased negative self-schema, potentially clarifying the confounded link between negative self-schema and psychosis proneness. Additionally, exploring the potential confounding role of depressive symptomatology in the causal relationship between social cohesion, trust, and psychosis proneness may offer insights into the impact of reduced active involvement in daily social interactions and the resulting perceived social isolation (Michalska da Rocha, Rhodes, Vasilopoulou, & Hutton, 2018). Secondly, focusing solely on the left-hemisphere narrowed our investigation to the precise neural networks accountable for language-related manipulation duties within the realm of WM. Thirdly, the graph illustrates complex relationships; for instance, a possibly direct effect on WM RT was observed for FEF and BA45, while no direct relationship between DLPFC and WM RT was noted. Future research could explore neural mechanism differences between individuals more prone to psychosis and those less prone, examining whether distinct pathways are engaged during working memory performance. Alternatively, investigations should explore whether DLPFC aberrations occur in individuals with psychosis proneness irrespective of the task used, without recruiting working memory-related brain regions. Further, detailed exploration of how these two mechanisms interact is warranted. Fourthly, the bootstrap analysis indicated low stability for the PAG (Supplementary Table 3). For example, the unconfounded effect of negative self-schema on PP is observed in 0.3016 out of 1000 bootstrapped samples, while in 0.3666 of the samples, such an unconfounded effect manifests in the opposite direction. Moreover, the unconfounded effect of SC&T on PP is detected in 0.1926 out of 1000 bootstrapped samples, surpassing the observations in the opposite and confounded directions. Based on our expert opinion and informed by prior studies, we posit that both negative self-schema and SC&T likely exert a direct influence on PP. When additional analysis was applied, incorporating negative self-schema and SC&T as direct causal effects of psychosis proneness, the effect sizes, RMSEA, and TLI values did not change (see Supplementary Table 6). Thus, the finding of possible latent variable by GFCI is not a drawback but rather a strength of the analysis which wouldn’t be possible with an expert opinion. Low stability is potentially due to the fact the data were sampled from the general population making effects related to psychotic experiences difficult to capture due to low population incidence of these experiences. This, in combination with a relatively small sample size and possible complex interaction among variables can all contribute to the observed low stability. Accounting for uncertainty allows meaningful conclusions and informed interpretations in line with the study’s goals. Future research can leverage these findings to refine causal discovery in a targeted population, enhance inferred relationships’ stability, by utilizing a dataset with larger sample size and advance our grasp of PP dynamics. Finally, in the current study, the scope was on examining the impact of individual risk factors on PP and the effects of PP on other variables. We did not investigate heritability in this study, and thus, we did not employ twin modeling. However, future studies can examine similar structures with twin modelling.

### Conclusion

To our knowledge, this is the first study to model direct causal pathways between environmental risk factors for psychosis, cognitive factors, negative schema of self, fMRI task-based activation during WM performance and, PP, using a data-driven CDA. The applied method, CDA, found the best fitting model based on computer calculations, as expert opinion would not have been optimal due to uncertainties regarding directional relationships between psychosis risk factors and other related factors. Our findings suggest the importance of targeting self-schema and SC&T in the general population as early as possible to prevent psychosis related experiences and possibly psychosis. These findings may alter perspectives on vulnerability to psychosis proneness. It appears that the internal environment, characterized by self-schema, or perception of the social environment hold greater importance than exposure to external risk events such as trauma. Furthermore, our findings suggest that although the effect was not strong compared to the population of patients, the functional brain, especially DLPFC, is affected by PP. These findings provide valuable insights into the underlying mechanisms of psychosis and have the potential to cover the path toward more effective interventions and treatments. If future research confirms the direct influence of negative self-schema and SC&T on psychosis proneness, screening for elevated levels of negative self-schema in the general population can be implemented to identify vulnerable individuals. Moreover, interventions targeting self-schema and SC&T deserve exploration regardless of confounding variables if the results are replicated. Tailored therapeutic interventions can then be selected for targeted treatment. Conducting screenings during adolescence or early adulthood may enhance the effectiveness of interventions before negative self-schema becomes established. Alternatively, online testing and a series of exercises can be developed to promote a more positive self-schema. For SC&T, activities fostering community integration and group inclusion can reduce the occurrence of psychosis proneness in the general population.

## Author contributions

Conceived and designed the study: TT. Collected data: TSI. Processed, Analyzed or Contributed to decisions about data analysis: TSI, SM, EK, ER, HYT, TT. Wrote the first draft of the paper: TSI. Contributed to and critically reviewed the manuscript: TT, SM, ER, EK, HYT.

## Acknowledgments.

We want to thank all the participants in this study. We also want to thank the following people who helped to collect data: Seda Arslan, Didenur Şahin-Çevik, Sara Sinem Sozan, Hande Ezgi Atmaca, Zeliha Oguz-Uguralp, Timuçin Baş or helped with aspects of the fMRI analysis: Zeliha Oguz-Uguralp and Kaan Uguralp. The following people helped review the manuscript: Fazilet Zeynep Yildirim-Keles, Elif Tugce Karoglu-Eravsar, Sumeyra Yalcintas and Daniel R Weinberger.

