## Supplementary for "Causal Discovery Analysis Reveals Insights into Psychosis Proneness, Brain Function, and Environmental Factors among Young Individuals"

**Participants**

**Exclusion and Inclusion Criteria**

Only if both members of the pair contributed MRI data and had less than 2 mm head movement in translation and rotation were included in the study. Exclusion criteria wereintellectual disability, head trauma, loss of consciousness, neurological diseases, any diagnosis of psychiatric diseases, using dental braces and medical conditions such as heart conditions, pregnancy. Participants whose age range between 14-24 were recruited for the study.

**Brief Core Schema Scales (BCSS)**

Items are rated on a five-point rating scale (0–4). Four dimensions of self and others, each with six items, are obtained: Negative Self (e.g. “I am unloved”), Positive Self (e.g. “I am respected”), Negative Others (e.g. “Other people are bad”), Positive Others (e.g. “Other people are good”).

**Environmental Assessments**

Ten measures of well-established environmental risk factors for psychosis were included. **Urbanicity.** Urbanicity was assessed using an amended version of the Medical Research Council sociodemographic schedule (1). Urbanicity was determined as the "place of birth". The level of urban/rural classification was determined through an open source "European Commission Global Human Settlement (GHS)" (<https://ghsl.jrc.ec.europa.eu/ESMVisualisation.php>). **Obstetric Complications.** Obstetric and delivery complications were evaluated using the ERI-RAOS risk factors questionnaire (2). **Cannabis Use.** Cannabis use was assessed as frequency, type, and duration of usage (Di Forti et al., 2009). **Childhood Trauma.** Childhood trauma was assessed by combining the information from the following scales: Childhood Trauma Questionnaire (CTQ) (3) and abuse rating questionnaires. Abuse questionnaires consisted of ratings of sexual abuse, physical abuse, psychological abuse, and bullying:

***Sexual Abuse Ratings Questionnaire***

Sexual Abuse Ratings Questionnaire assesses sexual acts towards a person, including intercourse, touching. During teenage years, willing contact with peers are excluded from being defined as sexual abuse. However, if any coercion or force is used, then it is included. Additively, non-contact verbal sexual solicitations are included if the perpetrator is a relative, known adult or an authority figure. Situations such that the subject is forced to watch sexual activity or pornography etc. Are also included. Since most abusers are adults, age difference between the subject and perpetrator was not conceived as a crucial factor. Sexual Abuse Ratings questionnaire includes indexes; the perpetrator (1 Both parents; 2 Mother; 3 Father; 4 Sibling; 5 Other relative; 6 Family friend; 7 Other), Frequency; Severity. The tool also includes ratings on "overall support received" and "official contact” (None; Social Services; General Practitioner; Police and other official contacts).

**Physical Abuse Ratings Questionnaire**

Physical abuse includes violent acts leading to physical injury or harm, such as harsh physical punishment etc. The questionnaire includes the same coding system as seen in sexual abuse ratings questionnaire for the perpetrator. The frequency, severity, support and official contact are also included as indexes.

***Psychological Abuse Ratings Questionnaire***

Psychological Abuse Ratings Questionnaire assesses experiences of humiliation, degradation such as being exposed to spreading lies about oneself by the perpetrator or shaming oneself in public or being exposed to terrorization such as invoking fear in the subject, in a calculated way. Furthermore, deprivation of basic needs such as light, sleep or lacking company of significant others, deprivation of social contact etc. Extreme rejection and emotional blackmail are also rated as psychological abuse.

***Bullying Ratings Questionnaire***

Bullying is an act of repetitively aggressive behavior by a peer with the intention to hurt the child, such as physical assault or intimidation or repeated name-calling). The Bullying Rating Questionnaire includes questions about teasing and bullying that participants may have experienced before 17 years of age by the perpetrator who similar age to participant is. Examples of incidents experienced by the perpetrator are such as; said mean and hurtful things or made fun of the participant or called subject mean and hurtful names; Completely ignored or excluded subject from their group of friends or left subject out of things on purpose; Hit, kicked or shoved the subject, or locked the subject in a room; Told lies or spread rumors about the subject; Other hurtful things.

**Parental Discord Ratings Scale.** Participants were asked about the level of argument and fighting in the family during their lifetime.

**Alcohol Usage.** Current alcohol use was measured based on the consumed alcohol more than 12 times in 12 months (4). **Discrimination.** The scale consists of 12 yes or no questions and participants were asked about the lifetime exposure to discrimination (5). **Life-Threatening Events.** Participants were asked about stressful life events in the last 12 months before the interview. The interview included 12 yes or no questions (6). **Social Cohesion and Trust (SC&T).** SC&T is one of the subdomains of the Social Environment Assessment Tool (SEAT)’s subdomains, and it looks at how people feel about being part of the community (Kirkbride, in preparation). Additionally, SC&T assesses whether people can be trusted and whether they are willing to help. A stronger sense of belonging and trust are indicated by higher SC&T scores.

**Categorization of Variables**

Before variables were introduced to Causal Discovery Analysis (CDA), variables were categorized as the following; for childhood trauma, all categories of questionnaires were collapsed and any severe level of trauma regardless of type and scale was regarded as exposure and counted as the existence of trauma (1) while no trauma (0) included participants with no exposure at all or exposures less than severe level; parental discord was divided into two categories; reporting of any level of discord (1), no report of discord (0); for alcohol usage, participants were divided based on the ones who consume alcohol more than 12 times in the last 12 months and participants who do not; for discrimination the total score was determined by summing up the yes answers. Summed score was used for the subsequent analysis; life threatening events total score was determined by summing up the yes answers; urbanicity were divided into two categories; : urban center (1) and suburb/rural areas (0); obstetric complications score were calculated into two categories; no complications (0), any complication (1); cannabis use were divided into two categories; no exposure in a lifetime (0), little/moderate exposure (little or a few times a year), and high exposure (a few times a month and above) (1) ; drug use other than cannabis were divided into two categories; no exposure in a lifetime (0), any exposure (1). For family history of psychosis, participants received a score of 0 in the absence of a family history of psychosis and a score of 1 if they had a first-degree family history of psychosis. Participants reported lifetime psychiatric medication use, receiving a score of 1 for users and 0 for non-users.

**Wechsler Abbreviated Scale of Intelligence | Second Edition (WASI-II).** Block Design measures the ability to analyze and synthesize abstract visual stimuli and matrix reasoning measures the capacities for fluid intelligence, broad visual intelligence, classification, and spatial ability, knowledge of part–whole relationships, simultaneous processing, and the perceptual organization of examinees. We converted the raw scores of the Block Design and Matrix Reasoning tests into age-adjusted scale scores. These scores were combined to calculate T scores. The sum of these T scores yielded the Perceptual Reasoning composite score.

**Verbal Fluency.** For phonemic fluency (letter fluency), words had to begin with a specified letter, such as a or s. For semantic fluency (category fluency), words had to belong to specified semantic categories, such as animals or vegetables.

**Neuroimaging**

**MRI Apparatus**

MR images were collected using a 3-Tesla MRI scanner (Siemens Magnetom Trio) with an echo-planar imaging (EPI) pulse sequence to generate anatomical images and blood oxygen level-dependent (BOLD) functional MRI data. 32- channel head coil was used for the application of radio-frequency (R.F.) pulses. Stimuli were presented by a 31.5" telemedicine LCD screen (1920 x 1080-pixel resolution and 59 Hz refresh rate) while participants were inside the scanner. The software from neurobehavioral systems (https://www.neurobs.com/) was used for stimulus presentation, reaction time (RT), and accuracy measurements.

**Structural Magnetic Resonance Imaging (sMRI)**

High-resolution T1-weighted images were acquired with TE=3.02 s; RT=2.6 s; Slice number =176; slice thickness = 1 mm; flip angle= 8°, field of view =256 mm; matrix=256 *256; voxel size= 1.0 x1.0x1.0 mm.

**Task-based Functional Magnetic Resonance Imaging (fMRI)**

Functional images were recorded with T2* weighted EPI with TR=2000 ms, TE=30 ms, tilt angle = 90 °, number of sections = 24, slice thickness: 5 mm; voxel size: 3.7 x 3.7 x 5.0 mm, FOV = 64x64 mm. All participants performed a WM task during the fMRI imaging.

**Working Memory Paradigm**

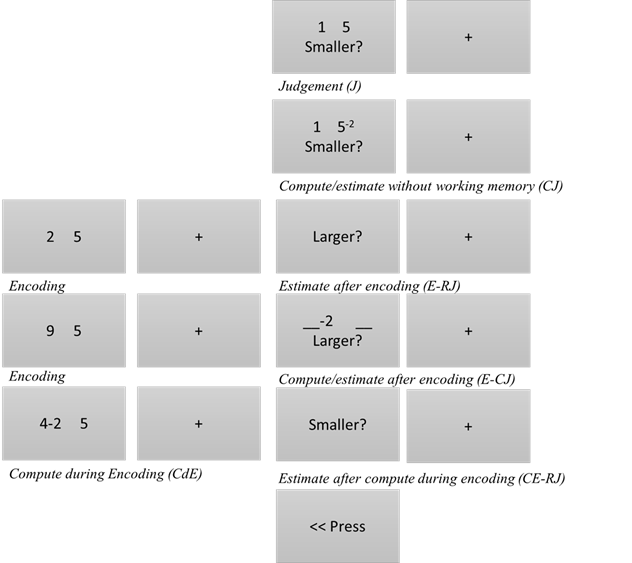

**Supplementary Figure 1.** *The Working Memory Paradigm*

Briefly, six different events comprised the paradigm ((Estimate without working Memory (J), Compute/estimate without working memory (CJ), Estimate after encoding (E-RJ), Compute/estimate after encoding (E-CJ) in which subjects performed subtraction on the retrieved number of the indicated side in working memory, Estimate after compute during encoding (CE-RJ), Control Motor Task (M) in which participants only asked to indicate the direction of an arrow (left or right), and each event has ten trials (Supplementary Fig 1). Two number digits were presented in the left and right side of the screen and participants were asked to respond which side of the number was larger or smaller. Depending on the specific event, participants were asked to either judge and respond without encoding (e.g., J, CJ) or with encoding (e.g., E-CJ, E-RJ) or without subtraction (e.g., J, E-RJ) or with subtraction (e.g., CJ, E-CJ, CE-RJ) on the retrieved number. The paradigm distinguishes well between encoding and response phases. The rest interval between encoding and response phases (the working memory interval) were jittered between 3 to 6 seconds and rest interval between each consecutive task were jittered between 10 seconds to 13 seconds.

**ROI Analysis**

Masks were created with MarsBar (7). Region of interest (ROI) analysis was conducted in Matlab 2020b. Each edge (ROI) constitutes the value derived from contrast estimates for each subject averaged over all the voxels in ROI during the manipulation phase of working memory paradigm. Average activation values during manipulation phase of the working-memory paradigm were extracted from 16 ROIs (Supplementary Table 1). We included ROIs from the left hemisphere and did not include the right hemisphere. Focusing on one hemisphere might neglect important interactions that occur across hemispheres. However, neuroimaging studies have demonstrated that certain aspects of working memory, such as verbal manipulation tasks, often exhibit lateralization towards the left hemisphere due to its language processing functions. Focusing solely on the left hemisphere in this investigation allowed us to narrow our scope to the specific neural networks responsible for linguistic manipulation tasks within the context of working memory while keeping the number of comparisons to a minimum. Time course normalization to raw fMRI data was applied with z-standardization to avoid signal changes between voxels due to physical and physiological aspects of MRI scan.

**Supplementary Table 1.**

|  | **Hemisphere** | **ROI Name-HCP** | **BA** | **MNI Coordinates** |
| --- | --- | --- | --- | --- |
| **1** | Left | Left BA 45 | BA 45 | -40 22 2 |
| **2** | Left | L 9-46d 173 |  | -33 50 13 |
| **3** | Left | L BA46 | BA46 | -34 38 30 |
| **4** | Left | BA6 (L 6a 42) | BA6 | -27 1 55 |
| **5** | Left | L SFL |  | -1 10 61 |
| **6** | Left | L AIP (BA40) | BA40 | -40 -46 42 |
| **7** | Left | L IP2 |  | -34 -53 43 |
| **8** | Left | Insula_L (L FOP4 72) |  | -42 12 0 |
| **9** | Left | BA9 (9a 174) | BA9 | -30 55 13 |
| **10** | Left | L-a9-46v (176) |  | -38 50 10 |
| **11** | Left | L-FOP4 |  | -36 16 8 |
|  |  | Frontal Operculum |  |  |
| **12** | Left | L-IFJa |  | -45 14 27 |
|  |  | Anterior Inferior frontal junction |  |  |
| **13** | Left | L-8C (172) | BA8 | -44 19 31 |
| **14** | Left | L-p47r (167) | BA47 | -44 44 4 |
| **15** | Left | L-a32pr (142) |  | -3 27 38 |
| **16** | Left | L-FEF (46) |  | -34 -3 56 |

**Adjusting Observed Variables for Family-Level Clustering using Linear Mixed Model**

In the current datasets, there are 194 participants from 97 families. There are on average 2 participants per family. To address the potential biases associated with family, a multilevel mixed-effects linear regression model was applied in Stata (StataCorp. 2021. Stata Statistical Software: Release 17. College Station, TX: StataCorp LLC) to subtract family-level unobservables (estimated random effects) from the original scores of environmental, psychological, and fMRI variables before they were introduced to CDA. Data were Z-standardized before this analysis. The residuals from the constant only model with a random intercept at the family level were included in the CDA analysis.

**Multilevel Mixed-Effects Linear Regression**

*Example for a constant-only model with a random intercept at the family level*;

e.g. for, mixed *depvar* || FamilyID:

which fits the following equation,

*depvar*_ij = b0 + V_j + e_ij

i - individual i;

j - family j;

V_j - Family-level unobservables;

e_ij - random noise;

The || FamilyID: notation implements a random intercept model at the level of families assuming that we have some information we do not observe at the family level that does not change within family across different individuals. And the random-effects model is fitted by -mixed-for a variance of this V_j.

**Causal Discovery Analysis with Greedy Fast Causal Inference (GFCI)**

While GFCI operates in a purely data-driven fashion, background knowledge (such as cognitive measures cannot cause a participant’s sex) can be entered (see first level below) to refine the data-driven fit. Our background knowledge included four levels. The first level included age, sex, urban birth, obstetric complications, family history of psychosis and paternal age. No other variable could cause these variables, and causation to each other in the first level was forbidden. The second level included education in years and socioeconomic status. The third level included parental discord. Participants were asked to indicate the earliest period they could remember parental discord, and these periods were temporally earlier than other factors. The remaining variables were included in the fourth level. No variable in the lower hierarchy could cause a variable in the higher hierarchy. While traditional power analysis is not applicable to CDA because null hypothesis testing is not typically applicable, the statistical power of CDA can be understood using simulated results (10). Accordingly, under the current sample size, CDA can achieve sufficient power to detect findings while limiting false positives.

Since GFCI considers the possibility of latent variables, the presented graph shows some uncertainty about the possibility of the latent variables. Our main results suggest the possible confounded direct relationship between Negative Self-Schema to Psychosis Proneness and SC&T and Psychosis Proneness. This possible confounded causal relationship (o-->) means the following: either negative self-schema is the direct cause of psychosis proneness, or there is a latent variable that is cause of negative self-schema and psychosis proneness or both. This can mean three possible causal graphs that are not distinguishable from our data using the algorithms we used, i.e.

(A) negative ->psychosis proneness

(B) negative <->psychosis proneness

(C) negative ->psychosis proneness and negative <->psychosis proneness

Based solely on the statistical relationships within the data, the algorithm cannot ascertain whether a direct relationship exists between negative self-schema and psychosis proneness. Without additional domain knowledge, we cannot definitively conclude the presence of a direct relationship.

**Structural Equation Modelling**

**Supplementary Table 2.** Structural Equation Modelling

| lhs | op | rhs | est | se | z | pvalue | ci.lower | ci.upper | Adjusted p-Value |
| --- | --- | --- | --- | --- | --- | --- | --- | --- | --- |
| p47r | ~ | BA9a-BA46v | 0.629858 | 0.051584 | 12.21039 | 0 | 0.528755 | 0.73096 | 0 |
| IFJa | ~ | BA8c | 0.696492 | 0.046341 | 15.02963 | 0 | 0.605665 | 0.787319 | 0 |
| BA9a-BA46v | ~ | BA9-BA46d | 0.835355 | 0.036407 | 22.94504 | 0 | 0.763999 | 0.906711 | 0 |
| IP2 | ~ | AIP | 0.739865 | 0.045778 | 16.16213 | 0 | 0.650142 | 0.829588 | 0 |
| BA9-BA46d | ~ | BA9a | 0.853831 | 0.037362 | 22.85288 | 0 | 0.780602 | 0.927059 | 0 |
| FEF | ~~ | BA6 | 0.464244 | 0.066149 | 7.018145 | 2.25E-12 | 0.334594 | 0.593894 | 2.136E-11 |
| Psychosis Proneness | ~~ | Negative Self | 0.536952 | 0.079848 | 6.724692 | 1.76E-11 | 0.380453 | 0.693451 | 1.43285E-10 |
| FOP4 | ~~ | BA45 | 0.413896 | 0.061926 | 6.683668 | 2.33E-11 | 0.292522 | 0.53527 | 1.66036E-10 |
| BA9a | ~ | BA8c | 0.429764 | 0.06477 | 6.635257 | 3.24E-11 | 0.302818 | 0.556711 | 2.05161E-10 |
| Life Threatening Events | ~ | Discrimination | 0.406565 | 0.064092 | 6.343418 | 2.25E-10 | 0.280946 | 0.532183 | 1.28092E-09 |
| Insula | ~~ | FOP4 | 0.335155 | 0.056816 | 5.89895 | 3.66E-09 | 0.223798 | 0.446513 | 1.89562E-08 |
| BA46 | ~ | Insula | 0.365787 | 0.06419 | 5.698503 | 1.21E-08 | 0.239977 | 0.491597 | 5.74104E-08 |
| SES | ~~ | Family History of Psychosis | 0.391677 | 0.071221 | 5.499426 | 3.81E-08 | 0.252086 | 0.531269 | 1.67067E-07 |
| BA8c | ~ | IP2 | 0.351474 | 0.064276 | 5.468187 | 4.55E-08 | 0.225495 | 0.477453 | 1.85113E-07 |
| BA6 | ~~ | a32pr | 0.237728 | 0.049479 | 4.804658 | 1.55E-06 | 0.140752 | 0.334704 | 5.89062E-06 |
| SFL | ~ | BA9-BA46d | 0.299678 | 0.06355 | 4.715652 | 2.41E-06 | 0.175123 | 0.424233 | 8.5834E-06 |
| SES | ~ | Education in Years | -0.26962 | 0.060443 | -4.46076 | 8.17E-06 | -0.38809 | -0.15116 | 2.73836E-05 |
| BA8c | ~ | SES | 0.26864 | 0.065135 | 4.124378 | 3.72E-05 | 0.140978 | 0.396301 | 0.000117717 |
| Cannabis Use | ~ | SES | -0.26867 | 0.067347 | -3.98942 | 6.62E-05 | -0.40067 | -0.13668 | 0.000198702 |
| p47r | ~ | IFJa | 0.203749 | 0.053302 | 3.822541 | 0.000132 | 0.099279 | 0.308218 | 0.000376438 |
| IFJa | ~~ | a32pr | 0.155949 | 0.041953 | 3.717227 | 0.000201 | 0.073723 | 0.238176 | 0.000546715 |
| Cannabis Use | ~ | Alcohol Usage | 0.243042 | 0.066065 | 3.678852 | 0.000234 | 0.113558 | 0.372526 | 0.000607015 |
| Insula | ~~ | a32pr | 0.203821 | 0.056338 | 3.617828 | 0.000297 | 0.093401 | 0.314242 | 0.000736256 |
| Education in Years | ~~ | Age | 0.233554 | 0.070927 | 3.292891 | 0.000992 | 0.09454 | 0.372569 | 0.002355115 |
| BA46 | ~~ | BA6 | 0.150623 | 0.046999 | 3.204783 | 0.001352 | 0.058506 | 0.24274 | 0.003081748 |
| BA9a-BA46v | ~ | Psychosis Proneness | 0.117197 | 0.036681 | 3.195032 | 0.001398 | 0.045303 | 0.18909 | 0.003065186 |
| Discrimination | ~ | IQ | 0.217828 | 0.068484 | 3.180688 | 0.001469 | 0.083601 | 0.352055 | 0.003101765 |
| Discrimination | ~ | Trauma | 0.205604 | 0.068564 | 2.998712 | 0.002711 | 0.071221 | 0.339987 | 0.005519298 |
| Psychosis Proneness | ~~ | Social Cohesion and Trust | -0.17872 | 0.061095 | -2.92531 | 0.003441 | -0.29847 | -0.05898 | 0.006763562 |
| AIP | ~~ | BA6 | 0.146315 | 0.050229 | 2.91299 | 0.00358 | 0.047869 | 0.244761 | 0.00680173 |
| Working Memory RT | ~~ | BA45 | -0.17103 | 0.059948 | -2.85295 | 0.004332 | -0.28852 | -0.05353 | 0.007964495 |
| Parental Discord | ~~ | Urbanicity | -0.20014 | 0.070877 | -2.82381 | 0.004746 | -0.33906 | -0.06123 | 0.008453178 |
| Cannabis Use | ~ | Working Memory RT | -0.18269 | 0.065523 | -2.78813 | 0.005301 | -0.31111 | -0.05426 | 0.009156854 |
| BA9a-BA46v | ~ | Psychiatric Medication | -0.10155 | 0.036542 | -2.77915 | 0.00545 | -0.17318 | -0.02993 | 0.009136886 |
| Education in Years | ~~ | Urbanicity | 0.186082 | 0.068476 | 2.717465 | 0.006578 | 0.051871 | 0.320294 | 0.010713414 |
| Trauma | ~ | Working Memory RT | 0.189537 | 0.069916 | 2.710906 | 0.00671 | 0.052503 | 0.326571 | 0.010624099 |
| Psychiatric Medication | ~ | SES | -0.19372 | 0.071803 | -2.6979 | 0.006978 | -0.33445 | -0.05299 | 0.01074976 |
| Working Memory RT | ~ | IQ | -0.18336 | 0.069 | -2.65732 | 0.007877 | -0.31859 | -0.04812 | 0.01181486 |
| Other Drug Use | ~ | BA9-BA46d | -0.18279 | 0.070375 | -2.59733 | 0.009395 | -0.32072 | -0.04485 | 0.013731557 |
| SFL | ~~ | FEF | 0.137109 | 0.053334 | 2.570771 | 0.010147 | 0.032577 | 0.241642 | 0.014459808 |
| Alcohol Usage | ~~ | Gender | -0.18365 | 0.071683 | -2.56198 | 0.010408 | -0.32415 | -0.04315 | 0.014469435 |
| IQ | ~ | Psychosis Proneness | 0.174152 | 0.070378 | 2.474539 | 0.013341 | 0.036215 | 0.31209 | 0.018105429 |
| SES | ~~ | OC | -0.1534 | 0.062008 | -2.47393 | 0.013364 | -0.27494 | -0.03187 | 0.017714416 |
| IP2 | ~ | Parental Discord | -0.11009 | 0.045735 | -2.40719 | 0.016076 | -0.19973 | -0.02045 | 0.020825716 |
| SFL | ~~ | a32pr | 0.135794 | 0.056784 | 2.391404 | 0.016784 | 0.024499 | 0.247089 | 0.021259823 |
| Life Threatening Events | ~~ | Verbal Fluency | 0.156615 | 0.065599 | 2.387458 | 0.016965 | 0.028043 | 0.285187 | 0.021022264 |
| IP2 | ~~ | Gender | -0.10893 | 0.046106 | -2.3626 | 0.018147 | -0.19929 | -0.01856 | 0.022008263 |
| FEF | ~~ | FOP4 | 0.089632 | 0.03815 | 2.349457 | 0.018801 | 0.014859 | 0.164405 | 0.022325978 |
| BA9-BA46d | ~ | Trauma | -0.08762 | 0.037323 | -2.34762 | 0.018894 | -0.16077 | -0.01447 | 0.021978382 |
| IP2 | ~~ | Family History of Psychosis | -0.09607 | 0.042155 | -2.27902 | 0.022666 | -0.17869 | -0.01345 | 0.025838926 |
| Life Threatening Events | ~~ | a32pr | -0.11822 | 0.054017 | -2.18856 | 0.028629 | -0.22409 | -0.01235 | 0.031996842 |
| AIP | ~~ | FOP4 | 0.101805 | 0.047601 | 2.138732 | 0.032457 | 0.008509 | 0.195101 | 0.035578316 |
| IQ | ~~ | Age | 0.139077 | 0.068788 | 2.021826 | 0.043194 | 0.004255 | 0.273899 | 0.046454332 |
| BA9a | ~~ | Urbanicity | 0.122682 | 0.062619 | 1.959191 | 0.05009 | -4.84E-05 | 0.245412 | 0.052873183 |
| Working Memory RT | ~~ | FEF | 0.104971 | 0.054202 | 1.936659 | 0.052787 | -0.00126 | 0.211206 | 0.054706595 |
| IFJa | ~ | Discrimination | 0.071138 | 0.046015 | 1.545963 | 0.122113 | -0.01905 | 0.161326 | 0.124294048 |
| p47r | ~~ | FOP4 | 0.048457 | 0.033268 | 1.456587 | 0.14523 | -0.01675 | 0.113661 | 0.145230378 |

**Partial Ancestral Graph (PAG) with Effect Sizes and p-values**

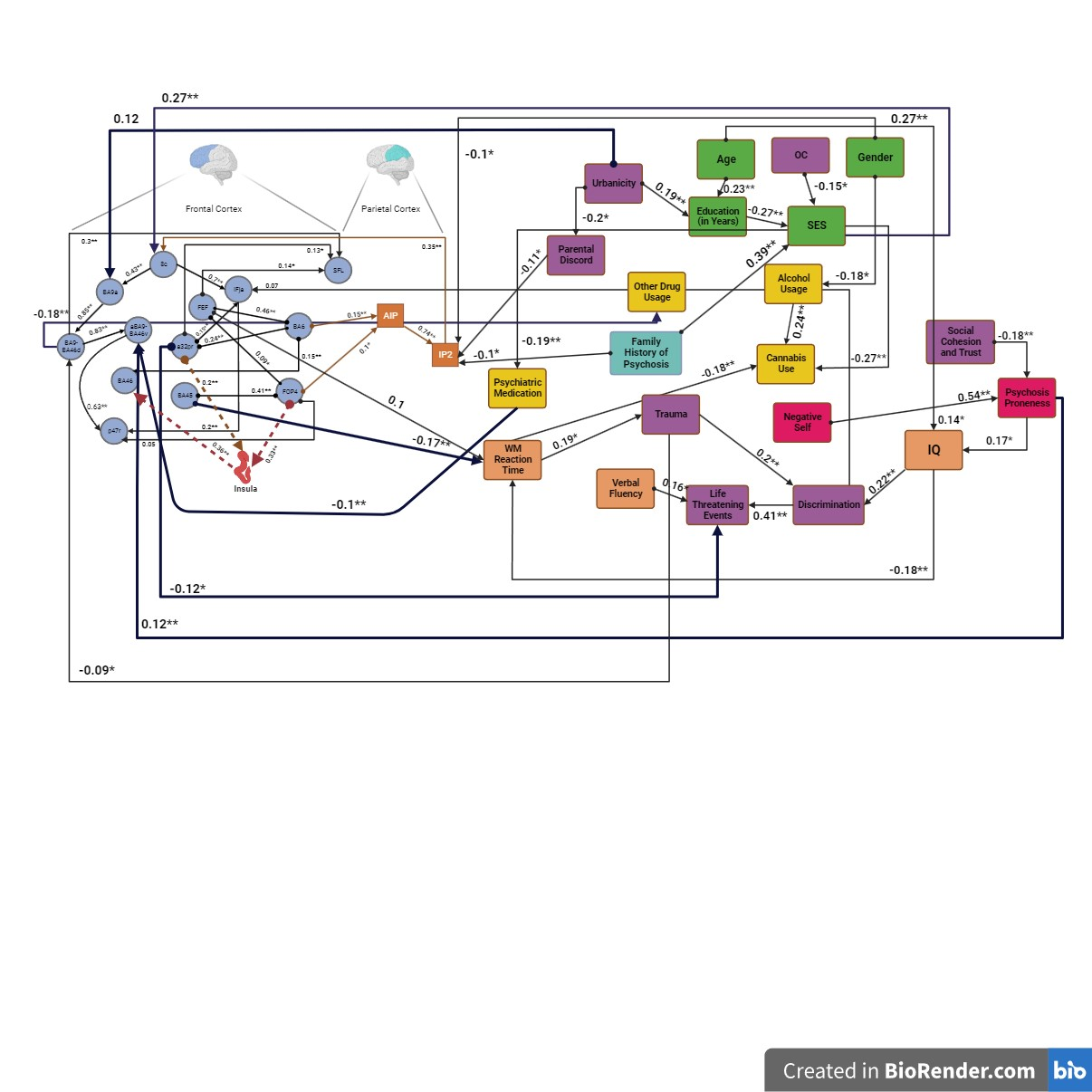

**Supplementary Figure 2. Complete Partial Ancestral Graph with Effect Sizes and p-value Information**

Causal Discovery Analysis was performed using GFCI for activations in ROIs when manipulating information during fMRI, environmental and psychological variables. (B.A.:Brodmann; SFL: Superior frontal language area; AIP: anterior intraparietal sulcus, I.P.: Intraparietal Sulcus; a9-46v: frontal mid, FOP4: frontal operculum, IFJa: anterior inferior frontal gyrus, p47r: the orbital prefrontal cortex, a32pr: refers to a narrow strip of the anterior midcingulate cortex, FEF: frontal eye fields; O.C.: obstetric complications; SES: socioeconomic status; W.M.: working memory)

** p< 0.01 * p< 0.5

(Created in BioRender.com)

**Re-Sampling Analysis**

**Supplementary Table 3.** Bootstrap Analysis

| **Bootstrap re-sampling analysis** | | | | | | | | | | | | | |
| --- | --- | --- | --- | --- | --- | --- | --- | --- | --- | --- | --- | --- | --- |
| **Edge Type in PAG** | **Nodes** | | **Proportion of 1,000 bootstrap resamples with edge type** | | | | | | | | | | |
|  | Node 1 | Node 2 |  |  |  |  |  |  |  |  |  |  | No Edge |
|  | Negative Self | Psychosis Proneness | 0 | 0 | 0.0186 | 0.007 | 0.3016 | 0.3666 | 0.1276 | 0.0812 | 0.0696 | 0.0186 | 0.0093 |
|  | Psychosis Proneness | aBA9-BA46v | 0 | 0 | 0.0162 | 0.0139 | 0.0696 | 0.0139 | 0.0162 | 0.0046 | 0 | 0 | 0.8654 |
|  | aBA9-BA46v | P47r | 0 | 0 | 0.0278 | 0 | 0.5661 | 0.2019 | 0.0835 | 0.0626 | 0.0557 | 0.0023 | 0 |
|  | IQ | WM Reaction Time | 0 | 0 | 0.0046 | 0.0209 | 0.0858 | 0.0882 | 0.0093 | 0.0232 | 0.007 | 0 | 0.761 |
|  | IQ | Discrimination | 0 | 0 | 0.0232 | 0.0139 | 0.1624 | 0.1137 | 0.0255 | 0.0186 | 0.0046 | 0.0046 | 0.6334 |
|  | WM Reaction Time | Trauma | 0 | 0 | 0.0186 | 0.0278 | 0.116 | 0.1323 | 0.0487 | 0.051 | 0.0278 | 0 | 0.5777 |
|  | BA45 | WM Reaction Time | 0 | 0 | 0.0116 | 0.0394 | 0.1647 | 0.1021 | 0.0162 | 0.0812 | 0.0046 | 0.007 | 0.5731 |
|  | Trauma | Discrimination | 0 | 0 | 0.0046 | 0.0162 | 0.116 | 0.1671 | 0.0371 | 0.0186 | 0.0162 | 0.0023 | 0.6218 |
|  | Discrimination | Life Threatening Events | 0 | 0 | 0.0302 | 0.0278 | 0.3248 | 0.3573 | 0.0742 | 0.1021 | 0.0418 | 0.007 | 0.0348 |
|  | Alcohol Usage | Cannabis Use | 0 | 0 | 0.0232 | 0.0278 | 0.2552 | 0.2297 | 0.058 | 0.0209 | 0.0186 | 0.0046 | 0.3619 |
|  | Psychiatric Medication | aBA9-BA46v | 0 | 0 | 0.0116 | 0.007 | 0.0742 | 0.0696 | 0.0232 | 0.0116 | 0.0046 | 0 | 0.7981 |
|  | Education in Years | SES | 0 | 0 | 0 | 0 | 0.29 | 0.239 | 0.0603 | 0 | 0 | 0.0209 | 0.3898 |
|  | Urbanicity | Education in Years | 0 | 0 | 0 | 0 | 0 | 0 | 0.3527 | 0 | 0 | 0.0302 | 0.6172 |
|  | Urbanicity | BA9a | 0 | 0 | 0 | 0 | 0.0023 | 0 | 0.1137 | 0 | 0 | 0 | 0.884 |
|  | SES | 8c | 0 | 0 | 0.0441 | 0 | 0.3271 | 0 | 0.0023 | 0 | 0 | 0.0023 | 0.6241 |
|  | SES | Cannabis | 0 | 0 | 0.0742 | 0 | 0.3202 | 0 | 0.0186 | 0 | 0 | 0.0974 | 0.4896 |
|  | Verbal Fluency | Life Threatening Events | 0 | 0 | 0.007 | 0 | 0.0858 | 0.051 | 0.0186 | 0.0116 | 0 | 0.0023 | 0.8237 |
|  | BA9a | BA9-BA46d | 0.0046 | 0 | 0.0162 | 0.0302 | 0.2854 | 0.4246 | 0.1392 | 0.0534 | 0.0441 | 0.0023 | 0 |
|  | FOP4 | BA45 | 0.0116 | 0.0023 | 0.0835 | 0.0093 | 0.522 | 0.1276 | 0.1206 | 0.0487 | 0.0534 | 0.0116 | 0.0093 |
|  | 8c | IFja | 0 | 0 | 0.0418 | 0.0116 | 0.5244 | 0.1833 | 0.1253 | 0.0418 | 0.051 | 0.0209 | 0 |
|  | BA9-BA46d | aBA9-BA46v | 0 | 0.0023 | 0.0139 | 0.0116 | 0.4478 | 0.2993 | 0.0696 | 0.1114 | 0.0302 | 0.0139 | 0 |
|  | BA6 | a32pr | 0 | 0 | 0.065 | 0.0139 | 0.3898 | 0.1763 | 0.1021 | 0.0487 | 0.0487 | 0.0186 | 0.1369 |
|  | AIP | IP2 | 0 | 0 | 0.0139 | 0.0162 | 0.4478 | 0.2923 | 0.1253 | 0.065 | 0.0209 | 0.0186 | 0 |
|  | BA6 | FEF | 0 | 0 | 0.0441 | 0.0093 | 0.3666 | 0.3039 | 0.1021 | 0.0812 | 0.0858 | 0.007 | 0 |
|  | Insula | BA46 | 0 | 0 | 0.0348 | 0.0139 | 0.297 | 0.2135 | 0.051 | 0.0278 | 0.0394 | 0.0023 | 0.3202 |
|  | FOP4 | Insula | 0.0139 | 0.0116 | 0.058 | 0.0441 | 0.3318 | 0.2854 | 0.0766 | 0.0673 | 0.0766 | 0.007 | 0.0278 |
|  | Family History of Psychosis | SES | 0 | 0 | 0 | 0 | 0 | 0 | 0.826 | 0 | 0 | 0.0302 | 0.1439 |
|  | 8c | BA9a | 0 | 0 | 0.0278 | 0.0209 | 0.0534 | 0.1276 | 0.0186 | 0.0394 | 0.0093 | 0.0139 | 0.6891 |
|  | FEF | FOP4 | 0 | 0 | 0.0093 | 0.0139 | 0.1114 | 0.0951 | 0.0232 | 0.0278 | 0.0116 | 0 | 0.7077 |
|  | IP2 | 8c | 0 | 0 | 0.007 | 0.0162 | 0.0464 | 0.0626 | 0.007 | 0.0139 | 0.0023 | 0 | 0.8445 |
|  | BA6 | AIP | 0 | 0 | 0.0255 | 0 | 0.1392 | 0.0626 | 0.0394 | 0.0093 | 0.0093 | 0.0046 | 0.71 |
|  | a32pr | IFJa | 0 | 0 | 0.0116 | 0.007 | 0.1253 | 0.1647 | 0.0232 | 0.0162 | 0.0093 | 0.0116 | 0.6311 |
|  | BA6 | BA46 | 0 | 0 | 0.109 | 0.0023 | 0.2042 | 0.0371 | 0.0603 | 0.0232 | 0.0093 | 0.0093 | 0.5452 |
|  | Psychosis Proneness | IQ | 0 | 0 | 0.0139 | 0.007 | 0.0858 | 0.0603 | 0.0139 | 0.0139 | 0.0046 | 0 | 0.8005 |
|  | Age | IQ | 0 | 0 | 0 | 0 | 0 | 0 | 0.2088 | 0 | 0 | 0.0023 | 0.7889 |
|  | BA9-BA46d | SFL | 0 | 0 | 0.0348 | 0.0139 | 0.2042 | 0.0742 | 0.0232 | 0.0209 | 0.007 | 0.007 | 0.6148 |
|  | FEF | SFL | 0 | 0 | 0.0719 | 0.0209 | 0.2622 | 0.0464 | 0.051 | 0.0255 | 0.0232 | 0 | 0.4988 |
|  | Age | Education in Years | 0 | 0 | 0 | 0 | 0 | 0 | 0.7146 | 0 | 0 | 0.0162 | 0.2691 |
|  | FOP4 | AIP | 0 | 0 | 0.0093 | 0.0046 | 0.0742 | 0.0464 | 0.0302 | 0.0232 | 0.0046 | 0.0116 | 0.7958 |
|  | FEF | Working Memory RT | 0 | 0 | 0.0325 | 0.0116 | 0.0603 | 0.0348 | 0.0325 | 0.0232 | 0 | 0 | 0.8051 |
|  | IFJa | P47r | 0 | 0 | 0.0209 | 0.0046 | 0.0557 | 0.0139 | 0.0046 | 0 | 0.0046 | 0 | 0.8956 |
|  | BA9-BA46d | Other Drug Usage | 0 | 0 | 0.0302 | 0.0046 | 0.1299 | 0.0186 | 0.0046 | 0.0278 | 0.0046 | 0 | 0.7796 |
|  | FOP4 | p47r | 0 | 0 | 0.0394 | 0.0209 | 0.116 | 0.0789 | 0.0418 | 0.0186 | 0.0046 | 0 | 0.6798 |
|  | OC | SES | 0 | 0 | 0 | 0 | 0 | 0 | 0.2877 | 0 | 0 | 0 | 0.7123 |
|  | a32pr | SFL | 0 | 0 | 0.0534 | 0.0162 | 0.174 | 0.0928 | 0.0464 | 0.0325 | 0.0116 | 0.0046 | 0.5684 |
|  | SC&T | Psychosis Proneness | 0 | 0 | 0.0116 | 0.0116 | 0.1926 | 0.1462 | 0.0928 | 0.0464 | 0.0186 | 0.007 | 0.4733 |
|  | SES | Psychiatric Medication | 0 | 0 | 0.0487 | 0 | 0.1671 | 0 | 0.007 | 0 | 0 | 0.0325 | 0.7448 |
|  | a32pr | LTE | 0 | 0 | 0.0116 | 0.0116 | 0.0696 | 0.0603 | 0.0023 | 0.0186 | 0.0023 | 0 | 0.8237 |
|  | Gender | Alcohol Usage | 0 | 0 | 0 | 0 | 0 | 0 | 0.3248 | 0 | 0 | 0.0093 | 0.6659 |
|  | Gender | IP2 | 0 | 0 | 0 | 0 | 0.0023 | 0 | 0.3202 | 0 | 0 | 0.0023 | 0.6752 |
|  | Trauma | BA9-BA46d | 0 | 0 | 0.0046 | 0.0023 | 0.0394 | 0 | 0.0232 | 0.0023 | 0 | 0 | 0.9281 |
|  | Discrimination | IFJa | 0 | 0 | 0.007 | 0.0023 | 0.0557 | 0.0278 | 0.007 | 0.0046 | 0 | 0 | 0.8956 |
|  | WM Reaction Time | Cannabis | 0 | 0 | 0.0139 | 0.0093 | 0.0951 | 0.0348 | 0.0418 | 0.007 | 0.0023 | 0 | 0.7958 |
|  | Parental Discord Severity | IP2 | 0 | 0 | 0.0302 | 0 | 0.0974 | 0 | 0.0673 | 0 | 0 | 0 | 0.8051 |
|  | Urbanicity | Parental Discord Severity | 0 | 0 | 0 | 0 | 0 | 0 | 0.3828 | 0 | 0 | 0.0302 | 0.587 |
|  | a32pr | Insula | 0 | 0 | 0.0302 | 0 | 0.0742 | 0.0209 | 0.0209 | 0.0046 | 0 | 0 | 0.8492 |
|  | Family History of Psychosis | IP2 | 0 | 0 | 0 | 0 | 0.0046 | 0 | 0.2158 | 0 | 0 | 0 | 0.7796 |

**SENSITIVITY ANALYSIS**

**Alternative Variable Coding of Environmental Risk Factors**

Since the level of effect of environmental risk factors on psychosis is still unknown, we conducted sensitivity analyses by coding environmental risk factors based on the odds ratios by (11). We coded the following environmental risk factors; urbanicity, obstetric complications, cannabis use, childhood adversity, paternal age and ethnic minority based on the (11). Briefly, urbanicity were divided into three categories; Urban center (1), Urban Cluster (0) and Rural (-1.5), obstetric complications score were calculated into two categories; no complications (0), complication with the largest OR score; Cannabis use were divided into three categories; no exposure (-1), little/moderate exposure (little or a few times a year) (0), and high exposure (a few times a month and above) (3); childhood trauma were divided into two categories; no childhood trauma (-1.5), existence of trauma (2.5), ethnic minority, native (0), others (2), paternal age; <40 (0), 40-50 (0.5), >50 (2). The remaining variables were coded in the same way as in the main analysis.

**Structural Equation Modelling of Sensitivity Analysis**

**Supplementary Table 4.** Structural Equation Modelling of the Sensitivity Analysis

| lhs | op | rhs | est | se | z | pvalue | ci.lower | ci.upper | Adjusted p-value |
| --- | --- | --- | --- | --- | --- | --- | --- | --- | --- |
| p47r | ~ | BA9a-BA46v | 0.629859 | 0.051581 | 12.21111 | 0 | 0.528762 | 0.730955 | 0 |
| IFJa | ~ | 8c | 0.696492 | 0.04634 | 15.0301 | 0 | 0.605668 | 0.787317 | 0 |
| BA9a-BA46v | ~ | BA9-BA46d | 0.835355 | 0.036401 | 22.94856 | 0 | 0.76401 | 0.9067 | 0 |
| IP2 | ~ | AIP | 0.740123 | 0.045743 | 16.18007 | 0 | 0.650469 | 0.829778 | 0 |
| BA9-BA46d | ~ | BA9a | 0.853831 | 0.037354 | 22.85773 | 0 | 0.780618 | 0.927043 | 0 |
| FEF | ~~ | BA6 | 0.464244 | 0.066149 | 7.018148 | 2.25E-12 | 0.334594 | 0.593894 | 2.06085E-11 |
| Psychosis Proneness | ~~ | Negative Self | 0.536946 | 0.079847 | 6.724673 | 1.76E-11 | 0.380449 | 0.693444 | 1.38276E-10 |
| FOP4 | ~~ | BA45 | 0.413895 | 0.061926 | 6.683654 | 2.33E-11 | 0.292521 | 0.535268 | 1.60227E-10 |
| BA9a | ~ | 8c | 0.43025 | 0.064704 | 6.649511 | 2.94E-11 | 0.303433 | 0.557068 | 1.79709E-10 |
| Life Threatening Events | ~ | Discrimination | 0.406564 | 0.064092 | 6.343417 | 2.25E-10 | 0.280946 | 0.532183 | 1.23598E-09 |
| Insula | ~~ | FOP4 | 0.335157 | 0.056816 | 5.898965 | 3.66E-09 | 0.223799 | 0.446515 | 1.82894E-08 |
| BA46 | ~ | Insula | 0.365787 | 0.06419 | 5.698517 | 1.21E-08 | 0.239977 | 0.491597 | 5.53913E-08 |
| SES | ~~ | Family History of Psychosis | 0.394167 | 0.071435 | 5.517808 | 3.43E-08 | 0.254156 | 0.534178 | 1.45223E-07 |
| 8c | ~ | IP2 | 0.351474 | 0.064264 | 5.469238 | 4.52E-08 | 0.225519 | 0.477429 | 1.77561E-07 |
| BA6 | ~~ | a32pr | 0.237727 | 0.049479 | 4.804646 | 1.55E-06 | 0.140751 | 0.334703 | 5.68427E-06 |
| SFL | ~ | BA9-BA46d | 0.299679 | 0.06354 | 4.716383 | 2.40E-06 | 0.175143 | 0.424215 | 8.25255E-06 |
| SES | ~ | Education in Years | -0.26594 | 0.060513 | -4.39484 | 1.11E-05 | -0.38455 | -0.14734 | 3.58641E-05 |
| 8c | ~ | SES | 0.26864 | 0.065129 | 4.124756 | 3.71E-05 | 0.14099 | 0.396289 | 0.0001134 |
| p47r | ~ | IFJa | 0.20375 | 0.053304 | 3.822414 | 0.000132 | 0.099276 | 0.308224 | 0.000382544 |
| IFJa | ~~ | a32pr | 0.155948 | 0.041953 | 3.717203 | 0.000201 | 0.073722 | 0.238175 | 0.000553962 |
| Insula | ~~ | a32pr | 0.203823 | 0.056338 | 3.617842 | 0.000297 | 0.093402 | 0.314244 | 0.000778039 |
| Education in Years | ~~ | Age | 0.235414 | 0.071025 | 3.314544 | 0.000918 | 0.096209 | 0.37462 | 0.002294821 |
| BA46 | ~~ | BA6 | 0.150624 | 0.046999 | 3.204802 | 0.001352 | 0.058507 | 0.242741 | 0.003231974 |
| BA9a-BA46v | ~ | Psychosis Proneness | 0.117197 | 0.036681 | 3.195021 | 0.001398 | 0.045303 | 0.18909 | 0.003204227 |
| Discrimination | ~ | IQ | 0.217828 | 0.068484 | 3.180723 | 0.001469 | 0.083602 | 0.352053 | 0.003231979 |
| Discrimination | ~ | Trauma | 0.205604 | 0.068564 | 2.998712 | 0.002711 | 0.071221 | 0.339987 | 0.005735299 |
| Psychosis Proneness | ~~ | Social Cohesion and Trust | -0.17872 | 0.061095 | -2.9253 | 0.003441 | -0.29846 | -0.05898 | 0.007009911 |
| AIP | ~~ | BA6 | 0.146315 | 0.050229 | 2.912983 | 0.00358 | 0.047869 | 0.244761 | 0.007032022 |
| WM Reaction Time | ~~ | BA45 | -0.17103 | 0.059948 | -2.85292 | 0.004332 | -0.28852 | -0.05353 | 0.008215692 |
| Parental Discord | ~~ | Urbanicity | -0.20151 | 0.070861 | -2.8438 | 0.004458 | -0.3404 | -0.06263 | 0.008172834 |
| BA9a-BA46v | ~ | Psychiatric Medication | -0.10155 | 0.036542 | -2.77916 | 0.00545 | -0.17318 | -0.02993 | 0.009669251 |
| Cannabis Use | ~ | Alcohol Usage | 0.191623 | 0.070465 | 2.71939 | 0.00654 | 0.053513 | 0.329732 | 0.011241051 |
| Trauma | ~ | WM Reaction Time | 0.189537 | 0.069916 | 2.710907 | 0.00671 | 0.052503 | 0.326571 | 0.01118323 |
| Psychiatric Medication | ~ | SES | -0.19372 | 0.071796 | -2.69814 | 0.006973 | -0.33443 | -0.053 | 0.011279419 |
| Education in Years | ~~ | Urbanicity | 0.183264 | 0.068342 | 2.681572 | 0.007328 | 0.049316 | 0.317212 | 0.011514967 |
| WM Reaction Time | ~ | IQ | -0.18336 | 0.068999 | -2.65735 | 0.007876 | -0.31859 | -0.04812 | 0.012032529 |
| Other Drug Usage | ~ | BA9-BA46d | -0.18279 | 0.070365 | -2.59772 | 0.009384 | -0.3207 | -0.04488 | 0.013949735 |
| SFL | ~~ | FEF | 0.137109 | 0.053334 | 2.570773 | 0.010147 | 0.032577 | 0.241642 | 0.014686716 |
| Alcohol Usage | ~~ | Gender | -0.18361 | 0.071679 | -2.5616 | 0.010419 | -0.3241 | -0.04312 | 0.014693804 |
| IQ | ~ | Psychosis Proneness | 0.174213 | 0.070379 | 2.475352 | 0.01331 | 0.036273 | 0.312153 | 0.018301906 |
| IP2 | ~ | Parental Discord | -0.10999 | 0.0457 | -2.40681 | 0.016093 | -0.19956 | -0.02042 | 0.021587706 |
| SFL | ~~ | a32pr | 0.135794 | 0.056784 | 2.391403 | 0.016784 | 0.024499 | 0.247089 | 0.021979223 |
| Life Threatening Events | ~~ | Verbal Fluency | 0.156615 | 0.065599 | 2.387456 | 0.016965 | 0.028043 | 0.285187 | 0.021699964 |
| IP2 | ~~ | Gender | -0.109 | 0.046074 | -2.36581 | 0.017991 | -0.19931 | -0.0187 | 0.02248843 |
| SES | ~~ | OC | -0.14593 | 0.061915 | -2.35694 | 0.018426 | -0.26728 | -0.02458 | 0.022520681 |
| FEF | ~~ | FOP4 | 0.089632 | 0.03815 | 2.349465 | 0.0188 | 0.01486 | 0.164405 | 0.022478734 |
| BA9-BA46d | ~ | Trauma | -0.08762 | 0.037323 | -2.34762 | 0.018894 | -0.16077 | -0.01447 | 0.022109635 |
| IP2 | ~~ | Family History of Psychosis | -0.09861 | 0.042175 | -2.33805 | 0.019384 | -0.18127 | -0.01595 | 0.022211405 |
| Life Threatening Events | ~~ | a32pr | -0.11822 | 0.054017 | -2.18854 | 0.02863 | -0.22409 | -0.01235 | 0.032136198 |
| AIP | ~~ | FOP4 | 0.101805 | 0.047601 | 2.138716 | 0.032459 | 0.008509 | 0.1951 | 0.035704562 |
| BA9a | ~~ | Urbanicity | 0.128457 | 0.062686 | 2.049218 | 0.040441 | 0.005595 | 0.251319 | 0.043612639 |
| IQ | ~~ | Age | 0.138916 | 0.068756 | 2.020431 | 0.043339 | 0.004157 | 0.273675 | 0.045839004 |
| WM Reaction Time | ~~ | FEF | 0.104972 | 0.054202 | 1.936674 | 0.052785 | -0.00126 | 0.211207 | 0.054777079 |
| IFJa | ~ | Discrimination | 0.071138 | 0.046015 | 1.545964 | 0.122113 | -0.01905 | 0.161326 | 0.124374713 |
| p47r | ~~ | FOP4 | 0.048457 | 0.033268 | 1.456572 | 0.145235 | -0.01675 | 0.11366 | 0.145234512 |

lhs: left-hand side, rhs: right-hand side

**Sensitivity Analysis- Graph with** **Effect Sizes and p-values**

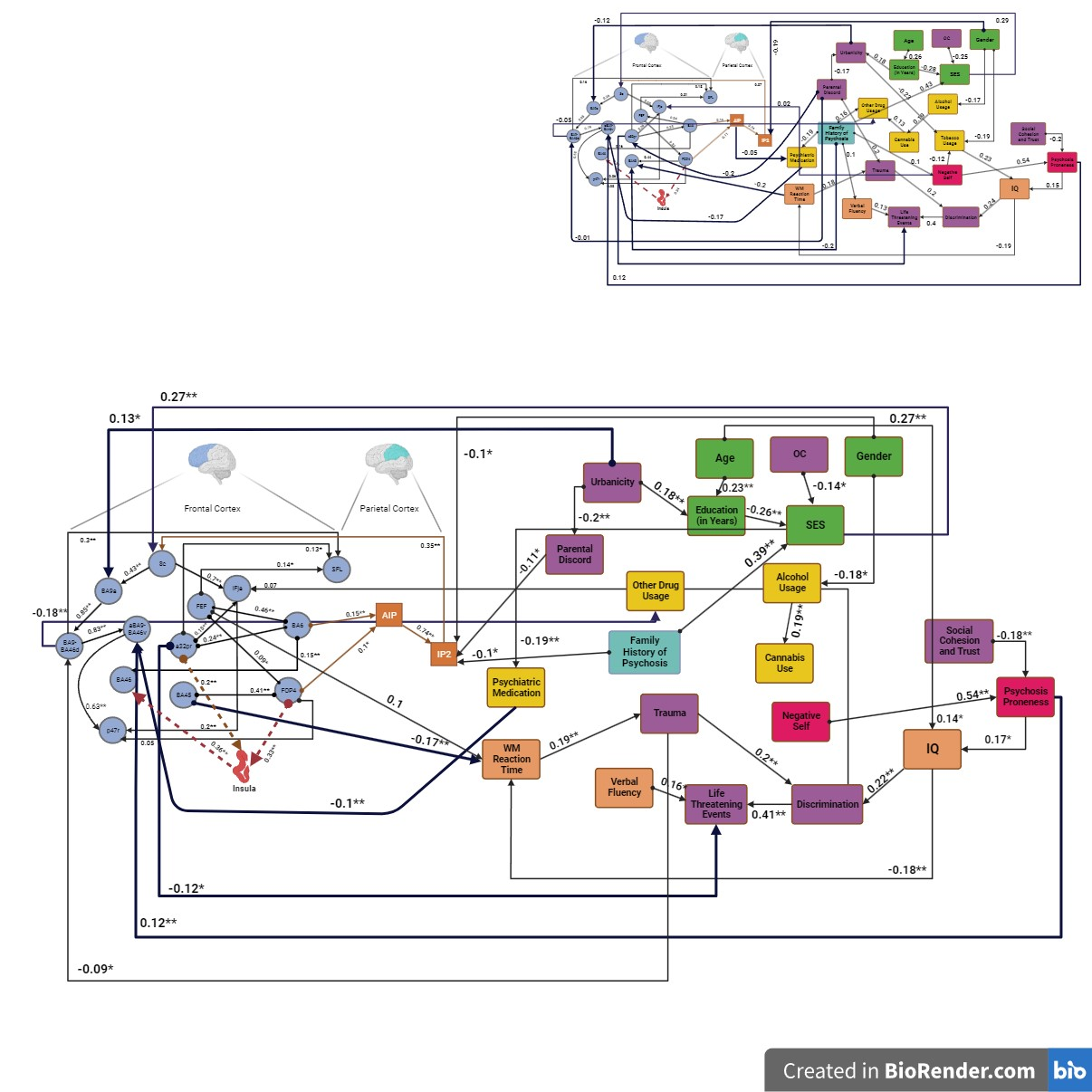

**Supplementary Figure 3. Sensitivity Analysis: Causal Discovery of Some Environmental Risk Factors Based on (11) and Multidomain Risk Factors and the Manipulation Phase of the Working Memory Paradigm**

Causal Discovery Analysis was performed using GFCI for activations in ROIs when manipulating information during fMRI, environmental risk factors based on the odds ratios by Vassos et al., 2020, other multidomain risk factors, and psychological variables. (B.A.:Brodmann; SFL: Superior frontal language area; AIP: anterior intraparietal sulcus, I.P.: Intraparietal Sulcus; a9-46v: frontal mid, FOP4: frontal operculum, IFJa: anterior inferior frontal gyrus, p47r: the orbital prefrontal cortex, a32pr: refers to a narrow strip of the anterior midcingulate cortex, FEF: frontal eye fields; O.C.: obstetric complications; SES: socioeconomic status; W.M.: working memory)

** p< 0.01 * p< 0.5

(Created in BioRender.com)

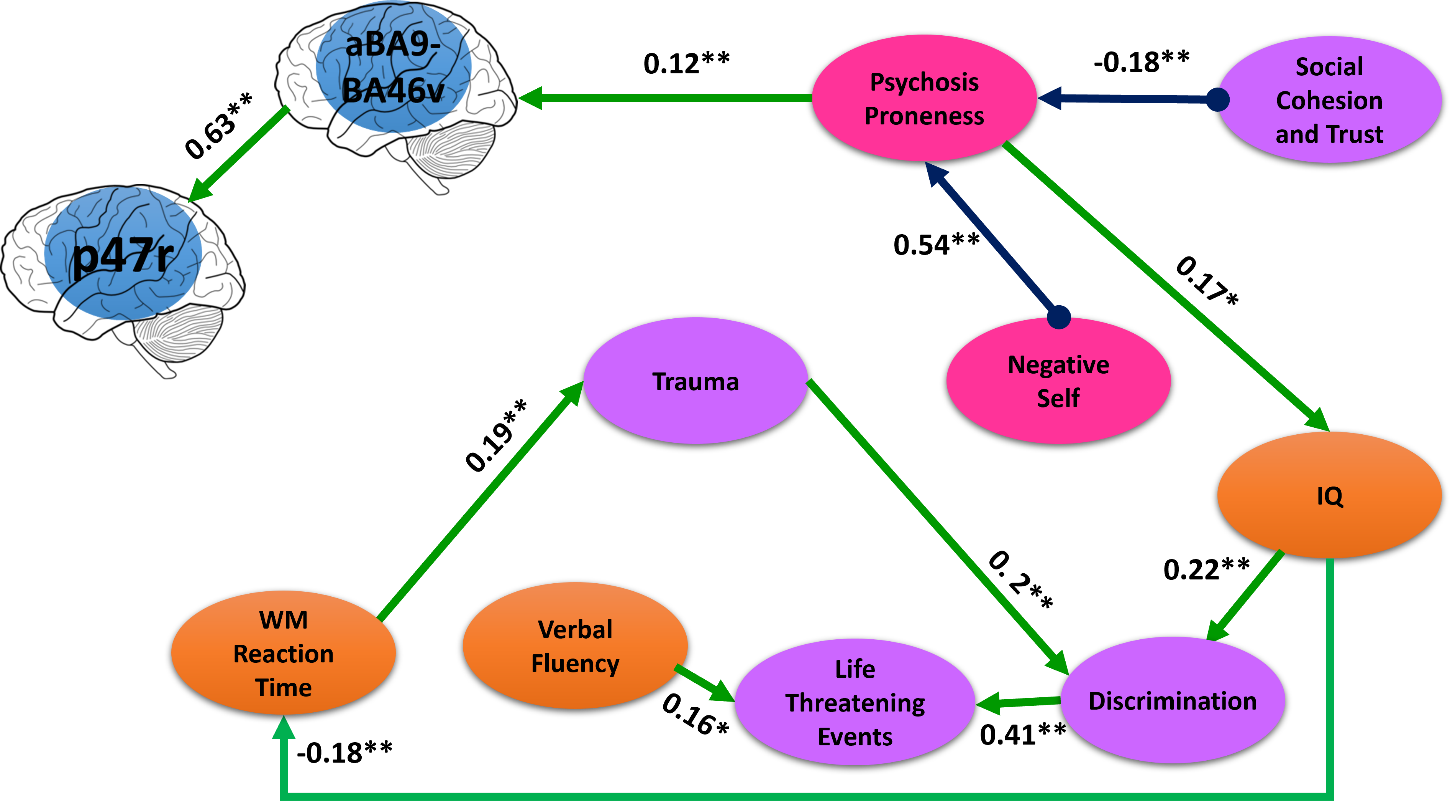

**Supplementary Figure 4. Psychosis Proneness Subgraph of the Sensitivity Analysis.**

GFCI returns a partial ancestral graph (PAG) depicting causal relationships between a set of variables, while assessing for unmeasured third variables in relationships (confounders). BA9: DLPFC; BA46: MPFC; P47r: Orbital Prefrontal Cortex

**Results of the Sensitivity Analysis and SEM Comparison with the Main Results**

Our sensitivity analysis, which incorporated risk factors including cannabis, trauma, urbanicity, obstetric complications, paternal age, and ethnic minority based on the odds ratio from (11), demonstrated that our primary findings remained unchanged. These findings continue to support the causal relationship between negative self-schema, SC&T and psychosis proneness, as well as the causal effect of psychosis proneness on aBA9-BA46v.

SEM results indicated no differences between alternative model and the original model. The Tucker-Lewis Index (TLI) and Root Mean Square Error (RMSEA) of both models are provided below.

|  | Main Model | Alternative model |
| --- | --- | --- |
| Tucker-Lewis Index | 0.833 | 0.831 |
| RMSEA | 0.052 | 0.052 |

In the alternative model in comparison to the main model, only two edges were removed. some edges were added, removed, and edge types were changed. The changed edges are provided below.

| **Node 1** | **Edge Type** | **Node 2** |
| --- | --- | --- |
| ***Removed Edges*** |  |  |
| WM Reaction Time | 🡪 | Cannabis Use |
| SES | 🡪 | Cannabis Use |

**Re-Sampling Analysis**

**Supplementary Table 5.** Bootstrap Analysis

| **Bootstrap re-sampling analysis** | | | | | | | | | | | | | |
| --- | --- | --- | --- | --- | --- | --- | --- | --- | --- | --- | --- | --- | --- |
| **Edge Type in PAG** | **Nodes** | | **Proportion of 1,000 bootstrap resamples with edge type** | | | | | | | | | | |
|  | Node 1 | Node 2 |  |  |  |  |  |  |  |  |  |  | No Edge |
|  | Negative Self | Psychosis Proneness | 0 | 0 | 0.0284 | 0.0071 | 0.3199 | 0.391 | 0.1209 | 0.0664 | 0.064 | 0.0024 | 0 |
|  | Psychosis Proneness | aBA9-BA46v | 0 | 0 | 0.0166 | 0.0118 | 0.0687 | 0.0284 | 0.0142 | 0.0071 | 0 | 0 | 0.8531 |
|  | aBA9-BA46v | P47r | 0 | 0 | 0.0355 | 0.0024 | 0.5806 | 0.2014 | 0.0616 | 0.0498 | 0.0616 | 0.0071 | 0 |
|  | IQ | WM Reaction Time | 0 | 0 | 0.0379 | 0.0095 | 0.1232 | 0.0948 | 0.0095 | 0.0403 | 0.0095 | 0.0047 | 0.6706 |
|  | IQ | Discrimination | 0 | 0 | 0.0474 | 0.0213 | 0.1588 | 0.1161 | 0.019 | 0.0213 | 0.0095 | 0.0047 | 0.6019 |
|  | WM Reaction Time | Trauma | 0 | 0 | 0.0024 | 0.0118 | 0.1043 | 0.2014 | 0.0308 | 0.0853 | 0.0142 | 0.0071 | 0.5427 |
|  | BA45 | WM Reaction Time | 0 | 0 | 0.0247 | 0.0261 | 0.2346 | 0.1422 | 0.0308 | 0.064 | 0.0047 | 0.0095 | 0.4645 |
|  | Trauma | Discrimination | 0 | 0 | 0.0047 | 0.0237 | 0.0782 | 0.1896 | 0.0427 | 0.0237 | 0.0118 | 0.0024 | 0.6232 |
|  | Discrimination | Life Threatening Events | 0 | 0 | 0.0498 | 0.0355 | 0.2796 | 0.4005 | 0.0782 | 0.0806 | 0.0379 | 0.0024 | 0.0355 |
|  | Alcohol Usage | Cannabis Use | 0 | 0 | 0.0284 | 0.0095 | 0.2038 | 0.1114 | 0.0521 | 0.0379 | 0.0118 | 0 | 0.545 |
|  | Psychiatric Medication | aBA9-BA46v | 0 | 0 | 0.0166 | 0.0071 | 0.1066 | 0.0829 | 0.0261 | 0.0118 | 0.0024 | 0 | 0.7464 |
|  | Education in Years | SES | 0 | 0 | 0 | 0 | 0.3578 | 0.2512 | 0.0427 | 0 | 0 | 0.0332 | 0.3152 |
|  | Urbanicity | Education in Years | 0 | 0 | 0 | 0 | 0.0024 | 0 | 0.3175 | 0 | 0 | 0.0498 | 0.6303 |
|  | Urbanicity | BA9a | 0 | 0 | 0 | 0 | 0.0071 | 0 | 0.0687 | 0 | 0 | 0 | 0.9242 |
|  | SES | 8c | 0 | 0 | 0.0308 | 0 | 0.372 | 0 | 0 | 0 | 0 | 0.0095 | 0.5877 |
|  | Verbal Fluency | Life Threatening Events | 0 | 0 | 0.0071 | 0.0024 | 0.0711 | 0.0616 | 0.0213 | 0.0071 | 0 | 0 | 0.8294 |
|  | BA9a | BA9-BA46d | 0.0071 | 0.0095 | 0.0118 | 0.0261 | 0.2559 | 0.4265 | 0.1398 | 0.0687 | 0.0427 | 0.0118 | 0 |
|  | FOP4 | BA45 | 0.0071 | 0.0047 | 0.0948 | 0.0118 | 0.545 | 0.1019 | 0.1185 | 0.0474 | 0.0521 | 0.0142 | 0.0024 |
|  | 8c | IFja | 0 | 0 | 0.0616 | 0.0071 | 0.5284 | 0.1862 | 0.1209 | 0.045 | 0.045 | 0.0237 | 0 |
|  | BA9-BA46d | aBA9-BA46v | 0 | 0 | 0.0213 | 0.0308 | 0.4408 | 0.2938 | 0.0853 | 0.0829 | 0.0427 | 0.0024 | 0 |
|  | BA6 | a32pr | 0 | 0 | 0.0545 | 0.0261 | 0.391 | 0.1611 | 0.1019 | 0.045 | 0.0521 | 0.0237 | 0.1445 |
|  | AIP | IP2 | 0 | 0 | 0.0261 | 0.0095 | 0.3863 | 0.263 | 0.1493 | 0.0877 | 0.0521 | 0.0261 | 0 |
|  | BA6 | FEF | 0 | 0 | 0.0474 | 0.0213 | 0.3768 | 0.2559 | 0.0853 | 0.0877 | 0.1066 | 0.0166 | 0.0024 |
|  | Insula | BA46 | 0 | 0 | 0.0427 | 0.019 | 0.3175 | 0.1682 | 0.0758 | 0.019 | 0.0379 | 0 | 0.3199 |
|  | FOP4 | Insula | 0.0047 | 0.0071 | 0.0498 | 0.0261 | 0.3815 | 0.2536 | 0.0948 | 0.0853 | 0.0569 | 0.0047 | 0.0355 |
|  | Family History of Psychosis | SES | 0 | 0 | 0 | 0 | 0 | 0 | 0.8294 | 0 | 0 | 0.0166 | 0.154 |
|  | 8c | BA9a | 0 | 0 | 0.0237 | 0.0213 | 0.0758 | 0.1137 | 0.0379 | 0.0521 | 0.0142 | 0.0142 | 0.6469 |
|  | FEF | FOP4 | 0 | 0 | 0.0095 | 0.019 | 0.1185 | 0.0782 | 0.0118 | 0.0213 | 0.0166 | 0.0024 | 0.7227 |
|  | IP2 | 8c | 0 | 0 | 0.0047 | 0.019 | 0.0427 | 0.0711 | 0.0095 | 0.0095 | 0.0095 | 0 | 0.8341 |
|  | BA6 | AIP | 0 | 0 | 0.0166 | 0.0047 | 0.1374 | 0.0829 | 0.0237 | 0.0095 | 0.0118 | 0.0047 | 0.7085 |
|  | a32pr | IFJa | 0 | 0 | 0.0142 | 0.0166 | 0.128 | 0.1919 | 0.0166 | 0.0355 | 0.0071 | 0.0308 | 0.5592 |
|  | BA6 | BA46 | 0 | 0 | 0.0829 | 0.0071 | 0.2038 | 0.045 | 0.0616 | 0.0095 | 0.019 | 0.0095 | 0.5616 |
|  | Psychosis Proneness | IQ | 0 | 0 | 0.0071 | 0.019 | 0.0758 | 0.0877 | 0.0166 | 0.0071 | 0 | 0.0024 | 0.7844 |
|  | Age | IQ | 0 | 0 | 0 | 0 | 0 | 0 | 0.2464 | 0 | 0 | 0.0047 | 0.7488 |
|  | BA9-BA46d | SFL | 0 | 0 | 0.0521 | 0.0095 | 0.1659 | 0.0687 | 0.0213 | 0.0308 | 0.0071 | 0.0024 | 0.6422 |
|  | FEF | SFL | 0 | 0 | 0.0711 | 0.0024 | 0.2275 | 0.0616 | 0.064 | 0.0284 | 0.0237 | 0.0047 | 0.5166 |
|  | Age | Education in Years | 0 | 0 | 0 | 0 | 0.0118 | 0 | 0.7488 | 0 | 0 | 0.0047 | 0.2346 |
|  | FOP4 | AIP | 0 | 0 | 0.019 | 0.0095 | 0.0877 | 0.0592 | 0.0166 | 0.0142 | 0.0071 | 0.0024 | 0.7844 |
|  | FEF | Working Memory RT | 0 | 0 | 0.0379 | 0.0047 | 0.0664 | 0.0474 | 0.0427 | 0.0261 | 0.0047 | 0.0024 | 0.7678 |
|  | IFJa | P47r | 0 | 0 | 0.0213 | 0 | 0.0664 | 0.0142 | 0.0024 | 0 | 0 | 0 | 0.8957 |
|  | BA9-BA46d | Other Drug Usage | 0 | 0 | 0.0142 | 0.0047 | 0.1327 | 0.0237 | 0.0071 | 0.0237 | 0.0118 | 0 | 0.782 |
|  | FOP4 | p47r | 0 | 0 | 0.0427 | 0.019 | 0.1209 | 0.0711 | 0.0403 | 0.0071 | 0 | 0 | 0.6991 |
|  | OC | SES | 0 | 0 | 0 | 0 | 0 | 0 | 0.3199 | 0 | 0 | 0.0024 | 0.6777 |
|  | a32pr | SFL | 0 | 0 | 0.0592 | 0.0142 | 0.2346 | 0.0687 | 0.0664 | 0.0521 | 0.0142 | 0.0095 | 0.481 |
|  | SC&T | Psychosis Proneness | 0 | 0 | 0.0213 | 0.0379 | 0.1682 | 0.1161 | 0.0995 | 0.0308 | 0.0308 | 0 | 0.4953 |
|  | SES | Psychiatric Medication | 0 | 0 | 0.0427 | 0 | 0.2062 | 0 | 0.0142 | 0 | 0 | 0.0261 | 0.7109 |
|  | a32pr | LTE | 0 | 0 | 0.019 | 0.0142 | 0.0664 | 0.0735 | 0.0142 | 0.0261 | 0.0095 | 0 | 0.7773 |
|  | Gender | Alcohol Usage | 0 | 0 | 0 | 0 | 0 | 0 | 0.3815 | 0 | 0 | 0.0024 | 0.6161 |
|  | Gender | IP2 | 0 | 0 | 0 | 0 | 0 | 0 | 0.3152 | 0 | 0 | 0 | 0.6848 |
|  | Trauma | BA9-BA46d | 0 | 0 | 0.0047 | 0 | 0.0521 | 0.0071 | 0.0237 | 0 | 0 | 0 | 0.9123 |
|  | Discrimination | IFJa | 0 | 0 | 0.0118 | 0.0071 | 0.0498 | 0.0498 | 0.0142 | 0.0118 | 0 | 0 | 0.8555 |
|  | Parental Discord Severity | IP2 | 0 | 0 | 0.0332 | 0 | 0.1019 | 0 | 0.0995 | 0 | 0 | 0.0024 | 0.763 |
|  | Urbanicity | Parental Discord Severity | 0.0024 | 0 | 0 | 0 | 0 | 0 | 0.391 | 0 | 0 | 0.0521 | 0.5545 |
|  | a32pr | Insula | 0 | 0 | 0.0071 | 0.0284 | 0.0853 | 0.0166 | 0.0142 | 0.0047 | 0 | 0 | 0.8436 |
|  | Family History of Psychosis | IP2 | 0 | 0 | 0.0024 | 0 | 0 | 0 | 0.1825 | 0 | 0 | 0 | 0.8152 |

**Supplementary Table 6.** Structural Equation Modelling with Negative Self-Schema and SC&T as a direct edge

| lhs | op | rhs | est | se | z | pvalue | ci.lower | ci.upper |
| --- | --- | --- | --- | --- | --- | --- | --- | --- |
| AIP | ~~ | FOP4 | 0.101806 | 0.047601 | 2.138745 | 0.032456 | 0.00851 | 0.195102 |
| AIP | ~~ | BA6 | 0.146315 | 0.050229 | 2.912993 | 0.00358 | 0.047869 | 0.244762 |
| Insula | ~~ | A32pr | 0.203821 | 0.056338 | 3.617826 | 0.000297 | 0.093401 | 0.314242 |
| BA46 | ~ | Insula | 0.365787 | 0.06419 | 5.698508 | 1.21E-08 | 0.239977 | 0.491597 |
| BA46 | ~~ | BA6 | 0.150624 | 0.046999 | 3.204811 | 0.001352 | 0.058507 | 0.242741 |
| FEF | ~~ | BA6 | 0.464243 | 0.066149 | 7.018136 | 2.25E-12 | 0.334593 | 0.593893 |
| SFL | ~ | BA9-BA46d | 0.299679 | 0.06355 | 4.715672 | 2.41E-06 | 0.175124 | 0.424234 |
| SFL | ~~ | FEF | 0.13711 | 0.053334 | 2.570789 | 0.010147 | 0.032578 | 0.241643 |
| SFL | ~~ | A32pr | 0.135795 | 0.056784 | 2.391418 | 0.016783 | 0.0245 | 0.24709 |
| FEF | ~~ | FOP4 | 0.089632 | 0.03815 | 2.349451 | 0.018801 | 0.014859 | 0.164405 |
| Insula | ~~ | FOP4 | 0.335157 | 0.056816 | 5.898961 | 3.66E-09 | 0.223799 | 0.446514 |
| WM RT | ~~ | BA45 | -0.17103 | 0.059948 | -2.85292 | 0.004332 | -0.28852 | -0.05353 |
| FOP4 | ~~ | BA45 | 0.413895 | 0.061926 | 6.683652 | 2.33E-11 | 0.292521 | 0.535269 |
| P47r | ~ | IFJa | 0.20375 | 0.053302 | 3.822569 | 0.000132 | 0.09928 | 0.308219 |
| P47r | ~ | BA9a-BA46v | 0.629859 | 0.051584 | 12.21042 | 0 | 0.528757 | 0.730961 |
| P47r | ~~ | FOP4 | 0.048457 | 0.033268 | 1.456588 | 0.14523 | -0.01675 | 0.113661 |
| Other drug use | ~ | BA9-BA46d | -0.18279 | 0.070376 | -2.59733 | 0.009395 | -0.32072 | -0.04485 |
| Discrimination | ~ | IQ | 0.217828 | 0.068484 | 3.180687 | 0.001469 | 0.083601 | 0.352055 |
| Discrimination | ~ | Trauma | 0.205604 | 0.068564 | 2.998712 | 0.002711 | 0.071221 | 0.339987 |
| IFJa | ~ | 8c | 0.696493 | 0.046341 | 15.02969 | 0 | 0.605666 | 0.78732 |
| IFJa | ~ | Discrimination | 0.07114 | 0.046015 | 1.545998 | 0.122105 | -0.01905 | 0.161328 |
| IFJa | ~~ | A32pr | 0.155948 | 0.041953 | 3.717205 | 0.000201 | 0.073722 | 0.238175 |
| BA6 | ~~ | A32pr | 0.237727 | 0.049479 | 4.804643 | 1.55E-06 | 0.140751 | 0.334704 |
| Life Threatening Events | ~ | Discrimination | 0.406564 | 0.064092 | 6.343418 | 2.25E-10 | 0.280946 | 0.532183 |
| Life Threatening Events | ~~ | Verbal Fluency | 0.156616 | 0.065599 | 2.387464 | 0.016965 | 0.028044 | 0.285188 |
| Life Threatening Events | ~~ | A32pr | -0.11822 | 0.054017 | -2.18856 | 0.028629 | -0.22409 | -0.01235 |
| Cannabis Use | ~ | Alcohol Usage | 0.243042 | 0.066065 | 3.67885 | 0.000234 | 0.113557 | 0.372526 |
| Cannabis Use | ~ | SES | -0.26867 | 0.067347 | -3.98943 | 6.62E-05 | -0.40067 | -0.13668 |
| Cannabis Use | ~ | WM RT | -0.18269 | 0.065523 | -2.78813 | 0.005301 | -0.31111 | -0.05426 |
| Education in Years | ~~ | Urbanicity | 0.186083 | 0.068476 | 2.717483 | 0.006578 | 0.051872 | 0.320295 |
| SES | ~ | Education in Years | -0.26962 | 0.060443 | -4.46074 | 8.17E-06 | -0.38809 | -0.15115 |
| Psychosis Proneness | ~ | Negative Self-Schema | 0.53973 | 0.05864 | 9.204106 | 0 | 0.424797 | 0.654662 |
| Psychosis Proneness | ~ | SC&T | -0.17965 | 0.05864 | -3.06356 | 0.002187 | -0.29458 | -0.06472 |
| WM RT | ~~ | FEF | 0.104971 | 0.054202 | 1.936656 | 0.052787 | -0.00126 | 0.211206 |
| WM RT | ~ | IQ | -0.18336 | 0.069 | -2.65732 | 0.007877 | -0.31859 | -0.04812 |
| Trauma | ~ | WM RT | 0.189537 | 0.069916 | 2.710907 | 0.00671 | 0.052503 | 0.326571 |
| 8c | ~ | SES | 0.26864 | 0.065134 | 4.124388 | 3.72E-05 | 0.140978 | 0.396301 |
| 8c | ~ | IP2 | 0.351474 | 0.064276 | 5.468184 | 4.55E-08 | 0.225495 | 0.477453 |
| BA9a-BA46v | ~ | Psychosis Proneness | 0.117197 | 0.036681 | 3.195022 | 0.001398 | 0.045303 | 0.18909 |
| BA9a-BA46v | ~ | BA9-BA46d | 0.835355 | 0.036407 | 22.94502 | 0 | 0.763999 | 0.906711 |
| BA9a-BA46v | ~ | Psychiatric Medication | -0.10155 | 0.036542 | -2.77915 | 0.00545 | -0.17318 | -0.02993 |
| BA9a | ~ | 8c | 0.429764 | 0.06477 | 6.63526 | 3.24E-11 | 0.302818 | 0.556711 |
| BA9a | ~~ | Urbanicity | 0.122683 | 0.062619 | 1.95921 | 0.050088 | -4.72E-05 | 0.245413 |
| IP2 | ~ | AIP | 0.739864 | 0.045778 | 16.1621 | 0 | 0.650142 | 0.829587 |
| IP2 | ~ | Parental Discord | -0.11009 | 0.045735 | -2.40715 | 0.016078 | -0.19973 | -0.02045 |
| BA9-BA46d | ~ | BA9a | 0.853831 | 0.037362 | 22.85289 | 0 | 0.780602 | 0.927059 |
| BA9-BA46d | ~ | Trauma | -0.08762 | 0.037323 | -2.34762 | 0.018894 | -0.16077 | -0.01447 |
| SES | ~~ | Family History of Psychosis | 0.391681 | 0.071222 | 5.499441 | 3.81E-08 | 0.252088 | 0.531273 |
| Parental Discord | ~~ | Urbanicity | -0.20014 | 0.070877 | -2.82382 | 0.004746 | -0.33906 | -0.06123 |
| Education in Years | ~~ | Age | 0.233555 | 0.070927 | 3.292908 | 0.000992 | 0.094541 | 0.372569 |
| Alcohol Usage | ~~ | Gender | -0.18365 | 0.071683 | -2.56197 | 0.010408 | -0.32415 | -0.04315 |
| SES | ~~ | OC | -0.1534 | 0.062008 | -2.47392 | 0.013364 | -0.27494 | -0.03187 |
| Psychiatric Medication | ~ | SES | -0.19372 | 0.071803 | -2.6979 | 0.006978 | -0.33445 | -0.05299 |
| IQ | ~ | Psychosis Proneness | 0.174152 | 0.070378 | 2.474532 | 0.013341 | 0.036214 | 0.31209 |
| IQ | ~~ | Age | 0.139077 | 0.068788 | 2.021834 | 0.043194 | 0.004256 | 0.273899 |
| IP2 | ~~ | Gender | -0.10893 | 0.046106 | -2.36261 | 0.018147 | -0.19929 | -0.01856 |
| IP2 | ~~ | Family History of Psychosis | -0.09607 | 0.042155 | -2.27902 | 0.022666 | -0.17869 | -0.01345 |

When additional analysis were conducted for the paths between Negative Self-Schema and PP, SC&T and PP, the effect sizes and general values of SEM did not change significantly. Rmsea: 0.051964; tli: 0.833262.
